# Efficient coding of natural images in the mouse visual cortex

**DOI:** 10.1101/2022.09.14.507893

**Authors:** Federico Bolaños, Javier G. Orlandi, Ryo Aoki, Akshay V. Jagadeesh, Justin L. Gardner, Andrea Benucci

## Abstract

How communication between neurons gives rise to natural vision remains a matter of intense investigation. The mid-level visual areas along the ventral stream, as studies in primates have shown, are selective to a common class of natural images—textures—but a circuit-level understanding of this selectivity and its link to perception remain unclear. We addressed these questions in mice, first showing that they can perceptually discriminate between texture types and statistically simpler spectrally matched stimuli. Then, at the neural level, we found that the secondary visual area (LM), more than the primary one (V1), was selective for the higher-order statistics of textures, both at the mesoscopic and single-cell levels. At the circuit level, textures were encoded in neural activity subspaces whose relative distances correlated with the statistical complexity of the images and with the mice’s ability to discriminate between them. These dependencies were more significant in LM, in which the texture-related subspaces were smaller and closer to each other, enabling better stimulus decoding in this area. Together, our results demonstrate texture vision in mice, finding a linking framework between stimulus statistics, neural representations, and perceptual sensitivity—a distinct hallmark of efficient coding computations.

## Introduction

Visual textures are broadly defined as “pictorial representations of spatial correlations” [1] —images of materials with orderly structures and characteristic statistical dependencies. They are pervasive in natural environments, playing a fundamental role in the perceptual segmentation of the visual scene [1, 2]. For example, textures can emphasize boundaries, curvatures [3, 4], 3D tilts and slants [5, 6] and distortions, support a rapid “pop-out” of stimulus features [7], and can form a basis set of visual features necessary for object vision [8].

Although texture images largely share the spectral complexity of other natural images [9–11], they can be more conveniently parametrized and synthetized than other natural images. This has been explored via diverse computational approaches: in the field of computer graphics [12], via entropy-based methods [13–15], using wavelet approaches [16, 17], and, more recently, in machine learning implementations based on deep convolutional neural networks [18–21].

In light of their rich statistics and convenient synthesis and parametrization, texture images have been at the core of studies on efficient coding principles of neural processing. According to the efficient coding hypothesis [22], the processing of visual signals along hierarchically organized cortical visual areas reflects the statistical characteristics of the visual inputs that these neural circuits have learned to encode, both developmentally and evolutionarily [23–29]. Accordingly, texture images have been extensively used in experimental studies that have examined the contribution of different visual areas to the processing of texture statistics.

In particular, studies in primates have revealed that the “mid-level” ventral areas, V2–V4, are crucial for processing texture images [30–41], more so than the primary visual cortex, V1 (however, see ref. [42]). Furthermore, as revealed by psychophysical observations [43] and neural measurements, area V2, in addition to being differentially modulated by the statistical dependencies of textures, correlates with the perceptual sensitivity for these stimuli [32, 40, 41]. Notably, biology-inspired computational studies using artificial neural networks have similarly emphasized hierarchical coding principles, with V2-like layers as the locus for representing texture images in classification tasks [44, 45]. Together, these observations suggest a general hierarchical coding framework, where the extrastriate visual areas, in particular area V2, define a neural substrate for representing texture stimuli, reflecting a progressive elaboration of visual information from “lower” to “higher” areas along the ventral visual stream.

This high-level view raises two fundamental questions: (1) whether this coding framework applies, in all generality, to hierarchically organized visual architectures as seen in several mammalian species other than primates—as CNN simulations would suggest—and (2) which functional principles at the circuit level give rise to texture selectivity, especially in the secondary visual area V2. Both questions hinge on the need to gain a computational and mechanistic understanding of how the visual system has evolved to process naturalistic statistical dependencies to enable the perception of scenes and objects [1, 2, 46–48].

Addressing these questions in the mouse model organism would be particularly advantageous [49]. Although the rodent visual system is much simpler than that of primates [50], mice and rats have a large secondary visual cortex (area LM) homologous to primate V2 [51, 52], belonging to a set of lateral visual areas forming a ventral stream of visual processing [53, 54]. As recordings from these areas have revealed, there is increased selectivity for complex stimulus statistics in both rats [55, 56] and mice [57, 58].

We studied the processing of texture images in mice with an emphasis on the interrelationship between behavioral, neural, and stimulus-statistic representations. Using a CNN-based algorithm for texture synthesis [59], we generated an arbitrary number of naturalistic texture exemplars and “scrambles”—spectrally matched images lacking the higher-order statistical complexity of textures [47, 60–63]—by precisely controlling the statistical properties of all the images. We demonstrated texture vision in mice showing they can perceptually detect higher order statistical dependencies in these natural images, distinguishing them from scrambles, and discriminating among the different types of naturalistic textures (“families” hereafter). At the neural level, using mesoscopic and two-photon GCaMP imaging, we found that the area LM was differentially modulated by texture statistics, more so than V1 and other higher visual areas (HVAs). Examining the representational geometry of the population responses, we found that when the statistical properties of a texture were most similar to those of scrambles, the corresponding neural activations were also more difficult to decode, and the animal’s performance decreased. These dependences were particularly prominent in LM and when considering the higher-order statistical properties of the images. Notably, LM encoded different texture families in neural subspaces that were closer to each other. Moreover, these subspaces were more compact in LM than in V1, thus enabling better stimulus decoding in this area.

## Results

### Training mice to detect and discriminate between texture statistics

To examine the ability of mice to use visual–texture information during perceptual behaviors, we designed two go/no go tasks. In the first task, the mice had to detect the texture images interleaved with scramble stimuli. In the second task, the mice had to discriminate between two types of texture images from different texture families

### Synthesis of textures and scrambles

We generated synthetic textures using an iterative model that uses a convolutional neural network (VGG16) to extract a compact multi-scale representation of texture images [59] (Figure 1a). To disentangle the contribution of higher-order image statistics from lower-order ones, for each texture exemplar we synthesized a spectrally matched image (scramble, Figure 1b) having the same mean luminance, contrast, average spatial frequency, and orientation content (Figure S1a-c, Methods) but lacking the higher-order statistical features characteristic of texture images. This produced image pairs for which the main axis of variation was higher-order statistics (textural information). In total, we synthesized images belonging to four texture families and four associated scramble families, each with 20 exemplars.

**Fig. 1:**
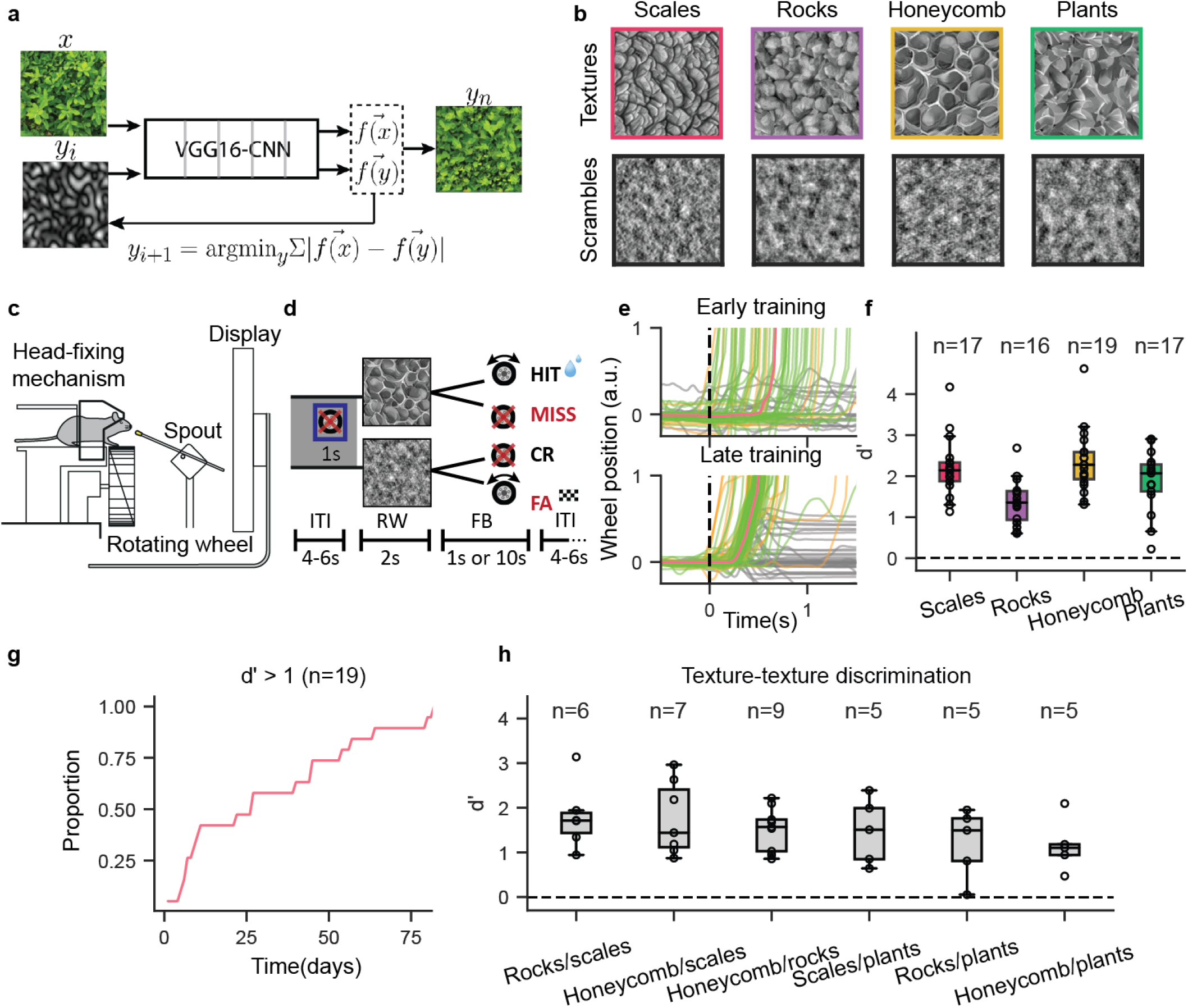
Mice can discriminate texture statistics from spectrally matched scrambles and between texture families. **a**, Schematic plot of the iterative algorithm to synthetize the texture images based on the VGG16-CNN architecture; ‘*x*’, target texture; ‘*f* (*x*)’, texture representation by the network; ‘*y*’ is an initial ‘seeding’ Gaussian-noise image with ‘*f* (*y*)’ being its network representation. The optimization minimizes the difference between *f* (*x*) and *f* (*y*) by iteratively changing ‘*y*_*i*_’ to obtain ‘*y*_*n*_’. **b**, Examples of texture families and the respective spectrally matched stimuli (scrambles). **c**, Schematic plot of the automatic training system with self-head fixation. **d**, The texture/scramble go no-go task: the mouse must rotate the rubber wheel (go trial) if shown a texture exemplar; it must keep it still if shown a scramble (no go). ITI is the inter-trial interval, RW the response window, and FB the feedback period. **e**, The representative examples of wheel rotations from an early training session (top) and a well-trained mouse (bottom); green for hits, yellow for false alarms, gray for either misses or correct rejects, and orange for the average across hits. **f**, Behavioral discriminability (*d*^′^) in the texture–scramble task for expert mice for each family. The top labels are the number of mice trained in each of the families; n = 16 out of 19 mice were trained in all the family-scramble pairs. The boxplots indicate the median and quartiles of the dataset; the black dots indicate the individual animals. Colors as in the image frames in (b). **g** The time needed for mice (proportion of days) to reach *d*^′^ ≥ 1 in their first family-scramble training. **h** Behavioral discriminability (*d*^′^) in the texture–texture task across all six possible pairs of the four families. The top labels are the number of mice trained in each texture pair. Each animal was trained in a different number of family pairs (Supplementary Table 1, 2).

### Behavioral detection of higher-order texture statistics

To train the mice in the two go/no go tasks, we employed an automated training setup [64], wherein the mice were asked to self-head fix and respond to the visual stimuli displayed on a computer screen located in front of them (Figure 1c). Mice were trained to respond to the target stimuli by rotating a toy wheel and, contingent on a correct response, they were rewarded with water. For the texture/scramble go/no go task, the “go” stimuli were texture images, while the “no go” stimuli were image scrambles (Figure 1d). For responses to a no-go stimulus (false alarms), a checkerboard pattern was displayed on the screen for 10 s before a new trial began. All the mice (n = 18) learned the task, with a *t*_50_ (i.e., the time needed for d-prime *>* 1 in at least half of the mice) being approximately 25 days (Figure 1g). Mice could significantly discriminate between all four texture/scramble pairs (Figure 1f, *d*^′^ *>* 1, *p <* 0.05 for all families, one sample t-test using the Holm-Bonferroni method to correct multiple comparisons; Supplementary Table 1) with an average discriminability value of *d*^′^ = 2.1 ± 0.15 (s.e.). The “rocks” family had a significantly lower performance than all other families but with a *d*^′^ still robustly larger than 1 (*d*^′^ = 1.4 ± 0.14, n = 16, *d*^′^ *>* 1, *p* = 0.016; *p* = 3 × 10^−5^, n = 15, ANOVA, repeated measures correction *p <* 0.03, n = 15, Tukey HSD). Dissecting the animals’ performance, we found that, on average, mice had a high proportion of hits (Figure S2a), as expected given that the training procedure encouraged “go” behaviors [65], with the lowest performance for rocks associated with a higher proportion of false alarms (Figure S2b). Additionally, to ensure that the mice were not adopting a strategy based on “brute force” memorization (e.g., of pixel-level luminance features [66]), we synthesized an additional 20 exemplars for each of the four families, together with corresponding scramble images. Then, in a subset of the mice (scales n = 4, rocks n = 3, honeycomb n = 11, plants n = 8), we switched the underlying set with the novel set and measured their performance over the last five sessions prior to the switch and the five sessions after the switch, finding no significant difference (Figure S2c).

### Behavioral discrimination between texture families

Having established that mice can detect higher-order statistical features in texture images that were missing in the scrambles, we examined whether they could discriminate between different texture statistics. We trained expert mice in texture–scramble discrimination, as well as a new cohort of naïve mice (n = 2), in a second go no-go task. They were shown exemplars (n = 20) from two texture families, randomly chosen but fixed across sessions, with only one of the two families associated with a water reward for a correct “go” response. In addition, all 40 exemplars were randomly rotated to prevent mice from solving this task using orientation information that may have been different across families (Figure S2d). Mice could discriminate between the texture families, with a significantly positive *d*^′^ for all six texture pairs (Figure 1h, *d*^′^ *>* 0, *p <* 0.02 for all pairs, one sample t-test with Holm-Bonferroni correction; Supplementary Table 2).

### Widefield responses to textures and scrambles

To examine the neural activations underlying the mice’s ability to detect and discriminate between texture statistics, we imaged multi-area responses from the posterior cortex of untrained animals whose neural dynamics were unaffected by procedural or perceptual learning processes. This choice assumes that texture processing in visual cortical networks is likely not the outcome of our behavioral training (see also Discussion).

We performed widefield calcium imaging during the passive viewing of textures and scrambles. Mice (n = 11) were placed in front of a computer screen that displayed either an exemplar of a texture or a scramble (Figure 2a). The stimuli, 100 degrees in size, were presented before the mice, centered on the mouse’s body midline, as was done for behavioral training. While the mice passively viewed the stimuli, we recorded both calcium-dependent and calcium-independent GCaMP responses using a dual wavelength imaging setup. We then used the calcium-independent GCaMP response to correct for the hemodynamic component of the calcium-dependent GCaMP responses [67]. We recorded from the right posterior cortex, which gave us access to ∼5–6 HVAs (Figure 2a). All the reliably segmented HVAs retinotopically represented the stimulus position in visual space (Figure S1d).

**Fig. 2:**
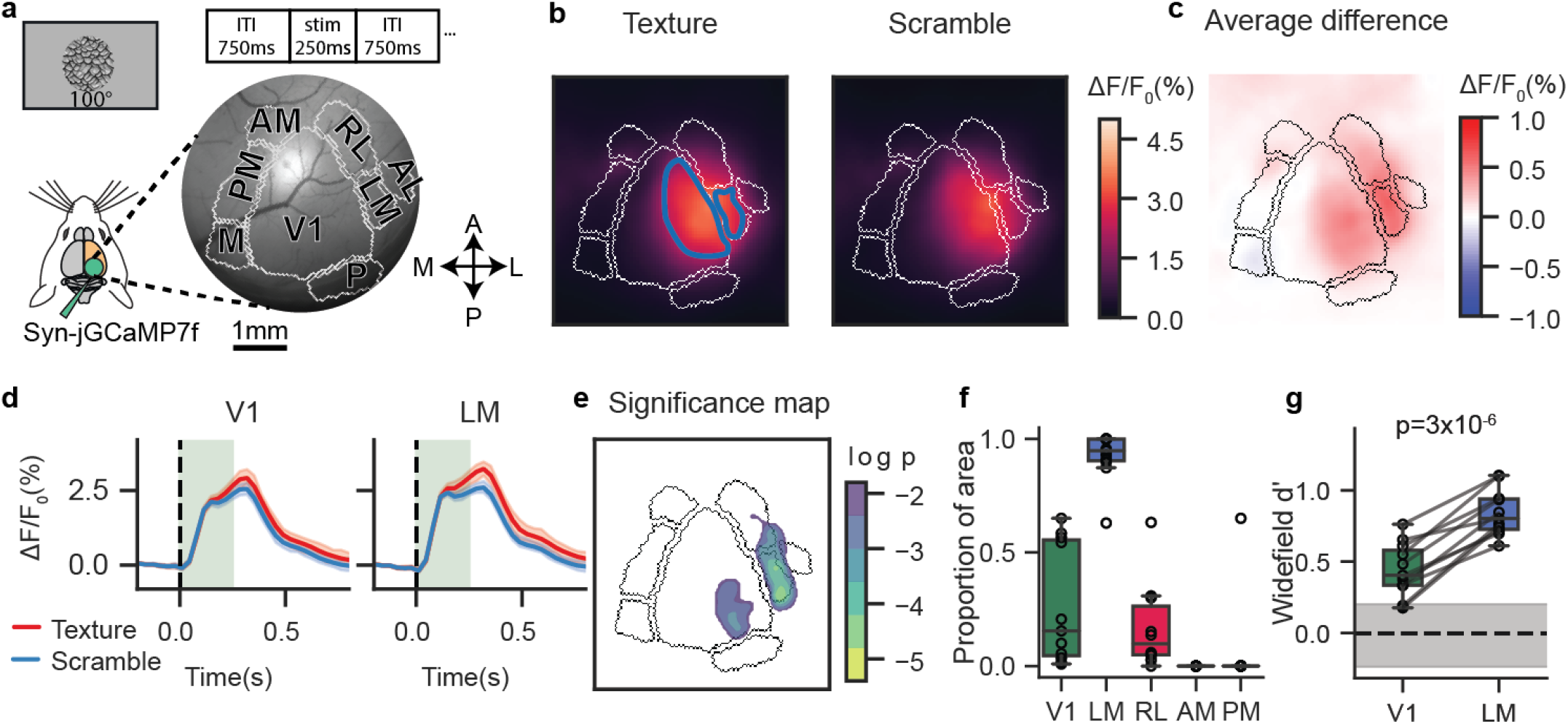
Texture stimuli differentially modulate V1 and LM at the mesoscale level. **a**, Schematic plot of the widefield imaging setup. Top, an example of a texture image and the stimulus presentation times. Right, a representative example of the right posterior cortex of a mouse with main-area borders (gray lines). V1, primary visual cortex; LM, secondary visual cortex (lateromedial); RL, rostrolateral; AL, anterolateral; AM, anteromedial; PM, posteromedial; M, medial area; P, posterior area. **b**, Average GCaMP response (across all exemplars and repeats) to texture stimuli (left) and scramble stimuli (right) for a representative example mouse. The blue contours show the regions of interest (ROIs) retinotopically matching the visual stimuli for both V1 and LM. **c**, The difference between the texture–scramble images shown in (b). **d**, The average GcaMP responses from the retinotopically matched ROIs in V1 and LM (Methods) for the same representative example shown in (b, c); textures in red, scrambles in blue. The green-shaded rectangles show the stimulus duration; the vertical broken line indicates the time of stimulus onset. **e**, The regions in V1 and LM with a statistically significant response difference shown in (c) – the logarithm of the p-values from a two-sided t-test; colored regions for p *<* 0.01. **f**, The proportion of pixels in each visual area (within the retinotopically identified ROIs) significantly modulated by the textures relative to the scrambles (n = 11 mice, empty circles). In color, V1, LM, RL having significantly positive values (p *<* 0.05, one sample t-test). **g**, The discriminability measure (*d*^′^) between the textures and scrambles from Δ*F/F* 0(%) responses within the same ROIs used for (f). The gray horizontal band corresponds to a null *d*^′^ distribution derived from pre-stimulus activity (Methods). The horizontal broken line indicates the mean of the null distribution; p-value from paired t-test, V1 vs LM *d*^′^ values (n = 10).

We computed the peak-response maps to the textures and scrambles showing activations almost exclusively in V1 and LM (Figure 2b). When averaging within the ROIs retinotopically matching the visual stimuli, the responses were larger for textures than scrambles both in V1 and LM (V1 scramble, average Δ*F/F* 0(%): 1.80% ± 0.11% s.e., V1 texture, average Δ*F/F* 0: 2.13% ± 0.11%, LM scramble Δ*F/F* 0(%): 1.63% ± 0.13%, LM texture Δ*F/F* 0(%): 2.14% ± 0.15%, n = 11). Similarly, the difference in the peak-response maps resulted in a differential modulation localized primarily in V1 and LM (Figure 2c,d, V1 Δ*F/F* 0(%) difference 0.27% ± 0.04%, LM Δ*F/F* 0(%) difference: 0.50% ± 0.03%). To establish statistical significance, we tested the modulation of each pixel against a null distribution derived from the pre-stimulus period (Figure 2e); and to determine the significance of an entire visual area, we computed the proportion of significantly modulated pixels in each area within retinotopic ROIs (Figure 2f). This analysis confirmed that the areas V1 and LM were those most significantly modulated by textures relative to scrambles (proportion of pixels in ROI *>* 0, V1: 0.27 ± 0.08 s.e. p = 0.007, LM: 0.92 ± 0.03 s.e. p = 6.4 × 10^−11^, RL: 0.17 ± 0.06 s.e. p = 0.021, AM: n.s., PM: 0.06 ± 0.06 s.e. p = 0.34; n = 11 mice, one sample t-test). Finally, to compare the V1 and LM modulations, we computed a texture discriminability measure (*d*^′^) in retinotopically matched ROIs and found that the *d*^′^ values in LM were significantly higher than those in V1 (Figure 2g. V1: 0.41 ± 0.05, s.e.; LM: 0.79 ± 0.05; difference, p = 3 × 10^−6^, paired t-test, n = 11). These results indicate that, at the mesoscopic level, when considering a constellation of HVAs surrounding the primary visual cortex, LM is the area with the most significant selectivity to higher-order texture statistics.

### Proportion of cells responding to textures and their modulation amplitude is higher in LM

We examined the circuit-level representations underlying this mesoscale selectivity using two-photon GCaMP recordings in areas V1 and LM (Figure 3a). Imaging ROIs (approximately 530 µm x 530 µm) in V1 and LM were selected based on the retinotopic coordinates of the visual stimuli, and neural activations were recorded while presenting three classes of visual stimuli: static gratings of different orientations and spatial frequencies (four orientations spaced every 45 degrees, 100 degrees in size, full contrast, sf = [0.02, 0.04, 0.1, 0.2, 0.5] cpd), scramble and texture images matching the properties of the stimuli used in behavioral experiments (four families for scrambles and textures, each with 20 exemplars rotated either by 0 or 90 degrees, and with eight repetitions of each image). The single-cell responses to oriented gratings agreed with what is typically reported in the literature (e.g. refs [68, 69]), with approximately 25–30% of the segmented cells being visually responsive (Figure 3c, V1 gratings: 25.37% ± 2.67% s.e., n = 6 mice; LM gratings: 27.71% ± 2.98%, n = 7; average no. of segmented cells = 381 ± 44 in V1 and 344 ± 46). The responses to textures and scrambles were rather heterogenous, with some cells strongly responding to textures, others to scrambles, and several showing mixed selectivity (Figure 3b). In both V1 and LM, there was a significantly larger proportion of cells responding to textures relative to gratings (Figure 3c, V1 textures: 61.05% ± 6.41% s.e., n = 6, LM textures: 55.27% ± 5.68% s.e., n = 7; gratings vs textures in V1, p = 0.0018, in LM, p = 0.015, paired t-test). Despite the significant heterogeneity, the responses, averaged across cells, were significantly larger in LM than in V1 for all texture families (Figure S3a,b; average V1 texture response: 9.5% ± 0.16%, s.e.; average LM texture response: 12.2% ± 0.18%, s.e.; *p <* 0.003 all families, paired t-test Holm-Bonferroni corrected). We then quantified the texture–scramble response modulation of the individual cells using a discriminability measure (*d*^′^), similar to what is done in mesoscale analyses (Figure 3d, e), and found that (i) the proportion of cells with significantly positive *d*^′^ values (i.e., with larger values in response to textures) were higher in LM than in V1 for all families (average proportion across families: V1: 67.5% ± 1.8% s.e., LM: 82.2% ± 1.4% s.e.. p-values for all families: scales = 7.4 × 10^−5^, rocks = 8.8 × 10^−4^, honeycomb = 0.024, plants = 0.006, paired t-test, Figure S4c); (ii) the average *d*^′^ value was higher in LM than V1 for all families (Figure 3f, Figure S4a,b, V1: average *d*^′^ = 0.24 ± 0.01, LM: average *d*^′^ = 0.54 ± 0.01, *p* = 2 × 10^−4^, paired t-test, n = 10), which reflected larger response amplitudes to textures than scrambles (Figure S4d, V1 texture/scramble difference: 1.5% ± 0.25, LM: 3.5% ± 0.39, *p* = 0.002, paired t-test, n = 10).

**Fig. 3:**
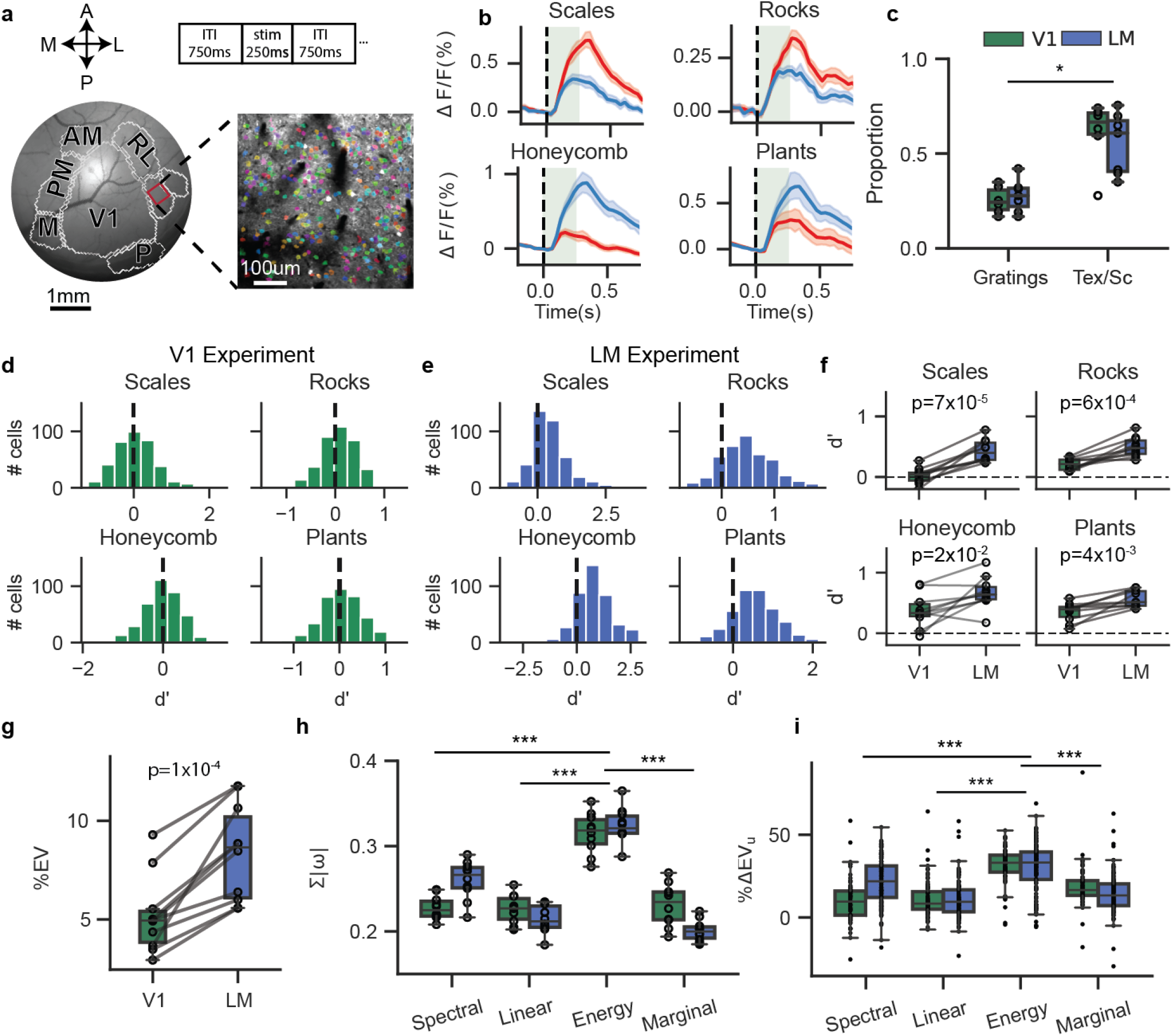
Single-cell responses in LM better discriminate textures from scrambles. **a**, Multi-area imaging, as in Figure 2a, with inset showing a representative ROI for two-photon recordings; colored dots indicate the segmented cells responsive to textures and/or scrambles (“stim”, top). **b**, Top panels: two example cells responding more strongly to a texture family (red) than scrambles (blue); bottom panels, two example cells for the opposite selectivity. **c**, The proportion of cells that significantly responded to oriented gratings and to either textures or scrambles (Txt/Scr) in V1 and LM; p-value, paired t-test across subset of animals (n = 6). **d, e**, The distributions of the texture–scramble discriminability values (*d*^′^) computed for each cell. Each panel is for a different texture family: green for V1; blue for LM. Data from a representative experiment. **f**, The mean *d*^′^ values for all the experiments in V1 and LM (n = 10 mice, black dots); connecting lines for the same-mouse data; V1 *d*^′^: scales = 0.02 ± 0.04 s.e., rocks = 0.21 ± 0.03, honeycomb = 0.37 ± 0.08, plants = 0.35 ± 0.05, LM *d*^′^: scales = 0.43 ± 0.05, rocks = 0.50 ± 0.05, honeycomb = 0.67 ± 0.08, plants = 0.56 ± 0.04; *p <* 0.001, obtained from paired t-tests with Holm-Bonferroni correction. **g** The explained variance (EV, %) by the encoding linear model based on PS image statistics, comparing V1 to LM; only cells for which EV ≥ 1% have been included in the analysis (permutation test, Methods); each empty dot is a mouse; connecting lines for the same-mouse data; p-value, paired t-test. **h**, The sum of weight values for each of the PS statistic groups of the fitted regressive model; each dot is an average across cells for a given mouse. The energy statistics are significantly higher than all others; one-way ANOVA with post-hoc analysis (Tukey HSD). Colors as in (g). **i**, The unique EVs for all four PS statistics groups. The cells with a high explained variance by the full model (EV ≥ 10%) were included in the analysis. Each dot is the change in explained variance for a cell when using the “full” model or a model missing a given PS statistic (Methods); the energy statistics are significantly higher than all others (one-ay ANOVA with post-hoc analysis, Tukey HSD). Colors as in (g).

Together, these results indicate that underlying the increased widefield texture selectivity in LM is both an increase in the proportion of texture-selective cells as well a larger texture–scramble modulation of individual cells.

### Encoding linear model of neural responses

To isolate the set of statistical features that most prominently drove the texture–scramble selectivity in V1 and LM, we used a previously described mathematical model to parametrize image statistics: the Portilla–Simoncelli statistical model (henceforth PS statistics [14]). This model employs a set of analytical equations to compute the correlations across a set of filters tuned to different image scales and orientations. These statistics can be divided into four main groups: marginal (skewness and kurtosis of the pixel histogram), spectral, linear cross-correlation, and energy cross-correlation statistics. The latter is best for distinguishing between texture and scramble images (Figure S6a, b). Using PS statistics as features, we created an encoding linear model for single-cell responses in V1 and LM. The model’s task was to predict the response of a particular neuron to all the texture and scramble exemplars as a weighted linear sum of PS coefficients. When considering the cells for which the model could explain at least 1% of the response variance—that is, a threshold value for the significance of the model’s fits derived from a permutation test (Methods)—we found that the proportion of these cells was higher in LM than in V1 (V1: 57.64% ± 3.77% s.e., LM: 78.20% ± 2.73%, n = 10), with a higher average explained variance in LM (Figure 3g, Figure S4e, V1: 5.27% ± 0.61% s.e., LM: 8.18% ± 0.67%, cross-validated, n = 10, V1/LM difference, *p* = 2 × 10^−4^, paired t-test). The energy cross-correlation statistics had the largest contribution to the explained variance (Figure 3h), which was also confirmed by an analysis of “unique” variance explained [70] (withholding a particular group of PS statistics, Methods), and it was found that the energy cross-correlation statistics was again the main contributor (Figure 3i).

As these results show, underlying the increased selectivity for textures in area LM and a larger proportion of cells having such selectivity is a stronger responsiveness to statistical features that are texture-defining—that is, those quantified by the energy cross-correlation PS statistics.

### Population responses to texture images

Next, we examined whether at the level of population encoding we could identify signatures of texture selectivity, more significantly so in LM than in V1. To discriminate the activity of the texture–scramble pairs separately in V1 and LM, we trained a binary logistic regression classifier. The decoder was largely above the chance level (50%) for all pairs (Figure 4a), with significantly larger performance in LM than in V1 when grouping all the texture families (p = 0.01, paired t-test). In both V1 and LM, the rocks family was the one with the lowest classification accuracy (Figure 4a, *p* = 4 × 10^−7^, n = 10, ANOVA; performance of rocks different from all pairs, repeated measures correction *p <* 0.035, n = 10, Tukey HSD). Notably, a similar drop in performance was also observed in the *d*^′^ measures of behavioral performance (Figure 4b,c, *p* = 3 × 10^−5^, n = 15, ANOVA, repeated measures correction *p <* 0.03, n = 15, Tukey HSD), where the lowest performance was observed for this texture–scramble pair consistently in individual mice, which were trained across all four texture–scramble pairs, and across animals.

**Fig. 4:**
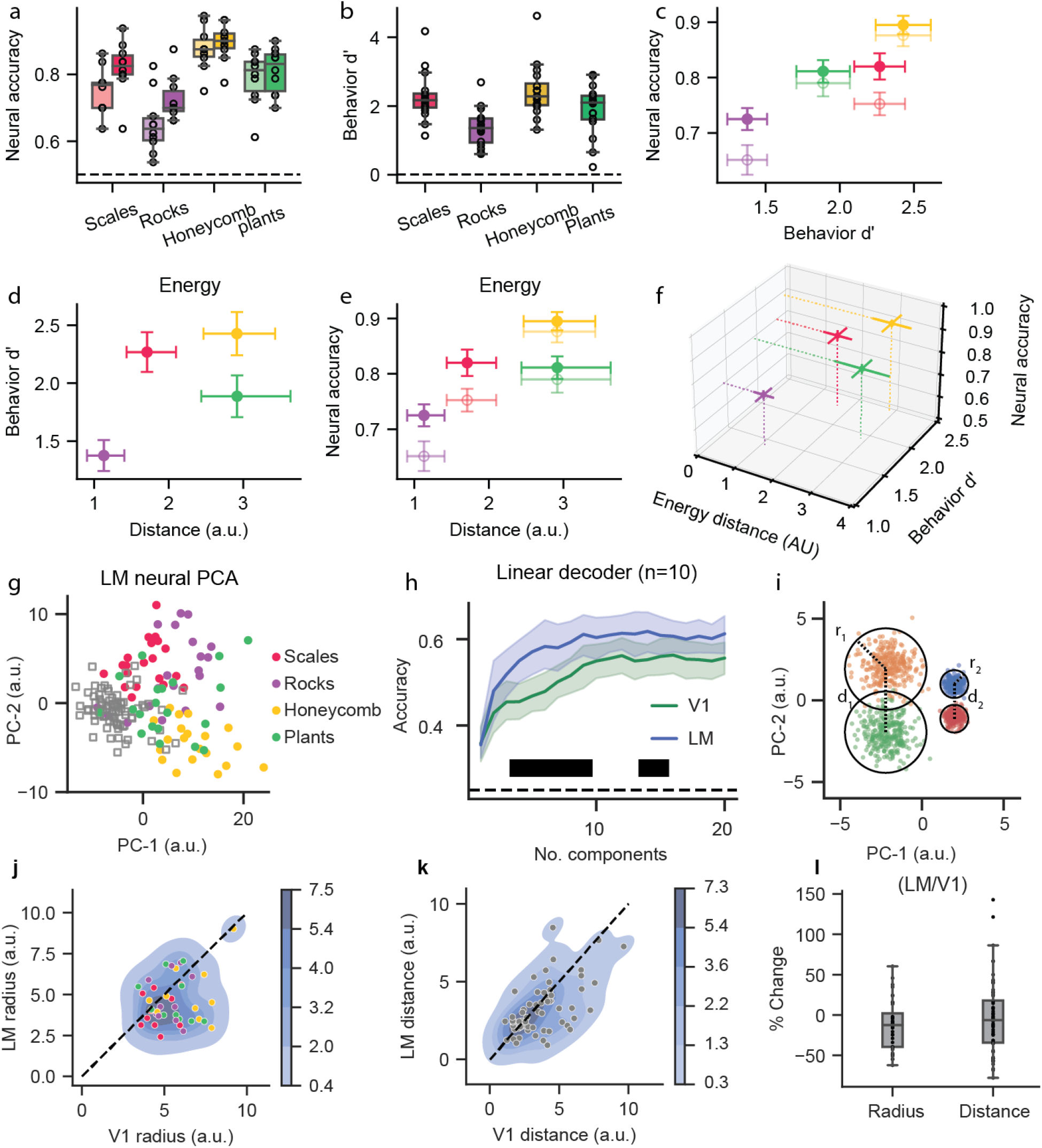
Statistical, behavioral, and neural discriminability correlate with the geometry of texture representations in LM. **a**, The accuracy of a linear classifier trained to discriminate textures from the scrambles for all four families using the neural responses from LM-ROI (saturated hue) and V1-ROI (low-saturation hue). The decoder accuracy is above 50% chance level for all pairs (V1: scales 75.2% ± 2.2% s.e. *p* = 1 × 10^−6^, V1: rocks 65.4% ± 2.8%, *p* = 3.5 × 10^−4^, V1: honeycomb 87.6% ± 2.1%. *p* = 2 × 10^−8^, V1: plant s 79.0% ± 2.5%. *p* = 1.1 × 10^−6^, LM: scales 82.2% ± 2.3%. *p* = 2 × 10^−7^, LM: rocks 72.5% ± 2.1%. *p* = 1.9 × 10^−6^, LM: honeycomb 89.5% ± 1.7%. *p* = 3 × 10^−9^, LM: plants 81.1% ± 2.1%. *p* = 1.5 × 10^−7^; n = 10 mice). **b**, The behavioral discriminability (*d*^′^), as in Figure 1f, but for a subset of the mice (n = 16) that completed the texture–scramble tasks for all four families. **c**, The combined 2D plot from the data in (a) and (b); the error bars are s.e.; color saturation as in (a). **d**, The behavioral discriminability as in (a) plotted against the inter-cluster distances for each texture/scramble family pair and for the energy cross-correlation statistics. The error bars are s.e. for the behavioral data and the 99.7% confidence intervals corrected for multiple comparisons (Šidák correction). **e**, The neural classifier accuracy, as in (b), against the inter-cluster distances for the energy cross-correlation statistics. Color saturation as in (a). **f**, The 3D plot of inter-cluster distances and the behavioral and neural discriminability measures; error bars as in (c-e). **g**, The 2D scatter plot for the first two PCA components of the neural responses from LM-ROI for one example animal; each dot is an exemplar (averaged across repeats, image rotations, and time frames around the peak response); filled circles for textures and empty squares for scrambles. **h**, The accuracy of a multinomial classifier (n = 10 mice) discriminating between the texture families as a function of the number of components, separately in V1 and LM PCA spaces. The shaded regions correspond to the 95% confidence intervals across all mice. The black bar indicates the range of PCA components for which the classifier accuracy is statistically different between V1 and LM (paired t-test, p-values *<* 0.05). **i**, The schematic illustrating the metrics measured in the neural PCA space. For every cloud of points in the PCA space, we measure its radius (e.g., r1, r2) and its distance with respect to another cloud (e.g., d1, d2). The clouds on the left show larger radii and inter-cluster distance compared to the clouds on the right. **j**, The scatter plot of the cluster radii in V1 (x-axis) and LM (y-axis) for all mice (n = 10). Each dot is a cluster radius for a given texture family and mouse. The shades of the blue contour lines are from a kernel-density estimation of the datapoints; the colors are as in (g). The black dotted line is the diagonal. **k**, Same as in (j) but for the inter-cluster distances. **l**, The decrease in radii and the inter-cluster distances in LM compared to V1 (% values relative to V1) for the data shown in (j,k); the difference is not significant; two-sided t-test.

### Linking image statistics to neural and behavioral representations

We next asked whether the correlation in neural and behavioral discriminability could be related to the statistics of the images. For instance, if, on average, the statistics of the rock exemplars were particularly similar to those of the scrambles compared to other families, then this reduced statistical discriminability may explain the drop in both behavioral and neural discriminability. We thus defined a distance metric in a statistical stimulus space based on a reduced set of PS statistics. In this space, a given visual stimulus is represented by a single point in four multi-dimensional (two dimensions per subspace) statistical subspaces for spectral, linear, energy, and marginal PS statistics (Figure S5a-d). In each subspace, we measured the inter-cluster (normalized) distances between the textures and the corresponding scrambles, finding that the rocks family had a significantly smaller texture–scramble distance than the other families in the energy statistics subspace (Figure 4d,e, x-axis, CI: 99.15%, multiple comparisons Šidák corrected CIs). For the other statistical subspaces, although the texture–scramble distances of rocks were still the shortest compared to the other families, they overlapped with at least one other family (Figure S7).

Together, the correlation between the PS-distance metric in the energy subspace, which best captures texture-defining statistics, and the drop in neural decoding and behavioral performance associated with the rocks family, suggest a tight linking framework between high-order image statistics, population encoding in V1 and LM, and behavioral performance (Figure 4f).

### V1 and LM differences in the representational geometry of texture families

The results from the binary logistic classifier trained to discriminate between the texture–scramble pairs suggest representational differences between V1 and LM. For instance, significantly fewer principal components (PCs) were needed in LM to attain maximum performance, two to four dimensions, whereas V1 required thrice as many, between four and 12 PCs (Figure S7; V1 accuracy, 12 dimensions: 78.6% ± 1.6% s.e.; LM accuracy, four dimensions: 78.6% ± 1.5%; *p <* 0.05 for mean accuracy of V1, 1-12 dimensions, vs LM, four dimensions, paired t-test, n = 10).

To examine the representational changes to texture stimuli between V1 and LM, we used a multinomial logistic regression classifier trained to categorize the four texture families across each of the 40 exemplar images for each family. Since the number of cells differed across experiments, we used PCA to fix the representational dimensionality of the activity space. Even with only two PCA components, the collective activations of the visually responsive cells across all texture and scramble stimuli already formed separate activity subspaces (clusters, Figure 4g) with an average explained variance above 15% (V1: 15.5% ± 1.4% s.e. LM: 19.1% ± 1.2%). The cross-validated classifier performed significantly above chance level in both areas, plateauing at approximately 60% performance with ∼10 PCA components (Figure 4h V1: 42.4% ± 3.0% s.e., accuracy *>* 25%, *p* = 2.3 × 10^−4^, LM: 48.0% ± 4.0%, accuracy *>* 25%, *p* = 2.6 × 10^−4^, one sample t-test). The LM decoder outperformed the V1 decoder, with significant differences observed reliably in the range between two and eight PCA components (Figure 4h). To highlight the properties of the population encoding that could explain the increased classification performance in LM, we studied the geometry of texture representations in a shared eight-dimensional PCA space of V1 and LM activations, in which the decoder had the largest significant discriminability power. Each point in this space corresponded to a texture exemplar (averaged across repeats) labeled according to the corresponding texture family (2D schematic of the 8D representations in Figure 4i,j). For every family, we computed the spread of the activations associated with the 40 exemplars—that is, the radii of the activity subspaces and their pairwise Euclidean distances (“inter-cluster” distances). In LM, we found that both the subspace radii and inter-cluster distances were significantly smaller than in V1 (Figure 4k,l; radii: p = 0.002, paired t-test, n = 10; inter-cluster distances: p = 0.02, paired t-test). The relative percentage decrease (V1 vs LM) was, on average, larger for the radii than for the inter-cluster distance values, but because of the large animal-to-animal variability, this difference did not attain statistical significance (Figure 4l, change in radius: −12.7% ± 4.7%, change in inter-cluster distance: −3.6% ± 5.7, LM relative to V1).

In conclusion, a population-level signature of the increased selectivity for energy cross-correlation statistics in individual cells in LM denotes a change in the representational geometry of the texture stimuli, with LM having more “compact” and better decodable representations than V1.

## Discussion

We found that mice can perceptually detect higher-order statistical dependences in texture images and discriminate between textures and scrambles and between different texture families. Across visual areas, V1 and LM were those most prominently modulated by texture statistics, with LM more so than V1, more significantly driven by the energy cross-correlation image statistics. The representational geometry of population responses demonstrated subspaces for each texture–scramble pair, with better stimulus decoding in LM than in V1. The distances between the texture–scramble subspaces changed according to the stimulus statistical dependencies, more significantly in the energy cross-correlation statistical components. The textures statistically most similar to scrambles (i.e., exemplars from the rocks family) had the shortest distances between the corresponding neural subspaces, with the worst perceptual discriminability by the animals and by a decoder trained on the neural representations. This was observed consistently in the animals trained on various texture–scramble pairs as well as across animals for this specific pair. Finally, the neural representations for different texture families were also easier to discriminate in LM than in V1, with LM having both shorter distances between texture subspaces as well as more compact subspaces (smaller radii) for individual textures, reflecting an overall more compact representational space for textures.

Efficiency, in reference to the efficient coding hypothesis [22], highlights a correspondence between input statistics, perceptual sensitivity, and the allocation of computational (and metabolic) resources. A neural code is efficient if it can reflect environmental statistics; such a code will favor basic visual features that are more common, relying on non-uniform neural representations and percentual sensitivity [23, 26–28]. This implies a close correspondence between neural, perceptual, and statistical representations. We studied this correspondence by examining the geometry of such representations in V1 and LM and identifying “rocks” as the family most similar to its scramble exemplars, with neural-distance representations and behavioral performance also being the smallest for this family. This was reliably observed in animals (tested across various texture–scramble pairs) and across animals for this pair. The selected texture families were chosen because of their likely ethological relevance to mice (e.g., rocks and plants) and their extensive use and characterization in the texture literature [40, 59]. They also had sufficiently diverse statistical dependences to permit a simple statistical similarity ranking between the texture–scramble pairs. However, future work could adopt a more principled approach in selecting texture families based on the statistical distance measure, as adopted in this study. This would allow us to define a psychometric difficulty axis in the stimulus-statistics space to be explored parametrically, both for texture–scramble and texture–texture discrimination. For the latter in particular, this approach could overcome a current statistical limitation: the six texture–family pairs span a relatively narrow range of distances in stimulus statistics, requiring an extremely large number of trials to test for differences in behavioral performance and neural representations, both within and across mice. Texture synthesis guided by a predetermined sampling of the relevant distances along a psychometric difficulty axis could ease the burden of collecting an exceedingly large dataset.

To examine the perceptual ability of mice to discriminate textures, we carefully controlled for the stimulus statistics of each exemplar. We customized a CNN-based approach for texture synthesis to achieve the equalization of lower (e.g., luminance, contrast, and marginal PS statistics) and higher statistical dependencies (e.g., linear and energy cross-features PS statistics). Further, we normalized the power spectrum in a frequency band of high perceptual sensitivity for mice and generated several metameric exemplars [43] differing in pixel-level representations but otherwise having identical statistical dependencies. We also introduced image rotations to ensure that the animals could generalize along this stimulus dimension. Finally, we tested the trained animals with new sets of metameric exemplars, confirming that “brute force” memorization of low-level features was not used in the task [66]. This approach gave us control over which statistical features the mice could use in the task and which component is critical when linking the statistical dependencies of the visual stimuli to neural and perceptual representations. In this respect, our approach may be preferable to using synthetic textures, in which typically only a reduced set of statistics of interest is under parametric control, while others are left free to (co)vary [13, 15, 18, 71, 72].

The prominent texture selectivity found in area LM is consistent with what is known about the area specialization of the mouse visual cortex, implicating LM in the processing of content-related (semantic) visual information [58, 69, 73–80], and with inactivation studies demonstrating the necessity of LM for the perception of even simple visual stimuli [74, 77].

At the circuit level, an analysis of the representational geometry of LM population responses [81, 82] revealed distinct activity subspaces associated with different texture families. These texture “manifolds” are reminiscent of the concept of object manifolds introduced in relation to the processing of complex objects along the ventral stream in primates [83–86] and in mice [57, 87]. When comparing LM to V1 representations, we found both a reduction in the size (diameter) of texture clusters and a reduction in inter-cluster distances. These two effects did not “compete” with each other in terms of signal decoding, leading to an overall improved linear discriminability of texture families in LM compared to V1. One interpretation is that the increased discriminability from V1 to LM is related to an increase in the representational invariances to low-level image statistics, as suggested by previous studies on rats [56] and mice [57]. The reduction in cluster sizes and the decrease in inter-cluster distances reflect an overall more compact representation of the four texture families, which may relate to LM achieving a higher encoding capacity than V1 while, at the same time, retaining large encoding accuracy for textures. Another possibility is that the V1 texture representations reflect a “partial inheritance” from LM via top-down signal processing [88]; experiments inactivating LM while recording from V1 could elucidate this point.

Neural recordings were done in untrained animals passively viewing the stimuli, thus enabling comparisons with primate studies that used similar preparations [32, 36, 37]. Furthermore, neural recordings in untrained animals eliminate the possibility that the observed selectivity and representational features emerge as a consequence of the task-learning process. Rather, our analyses likely highlight a computational property of the visual system emerging from an evolutionarily refined genetic program [29] and from exposure to a rich set of image statistics during development. The observation that in naïve animals the decoding quality of the neural signals follows the statistical separability of texture–scramble images, mirrored by congruent performance modulations in trained animals, supports this interpretation. It is also conceivable that learning and attentional processes, as animals engage in tasks, might affect the properties of neural representations [1, 89, 90]. Therefore, in future studies, it would be of interest to examine the neural dynamics underlying texture representations during the different phases of learning.

In conclusion, our results demonstrate the signal processing of naturalistic stimuli in the mouse visual cortex akin to what has been observed in primates, additionally highlighting an intimate link between the geometry of neural representations, stimulus statistical dependencies, and perceptual behavior, which is a distinct hallmark of efficient coding principles of information processing. Considering that similar processing features are also found in V2/LM equivalents in artificial neural networks, our results likely reflect a general efficient coding principle emerging in hierarchically organized computational architectures devoted to the extraction of semantic information from the visual scene.

## Acknowledgments

We thank D. Zoccolan for his feedback on the interpretation of our findings. We thank Yuki Goya, Rie Nishiyama, and Yuka Iwamoto for their support with behavioral training, animal care, and surgeries. We thank O’Hara and Co., Ltd., for their support with the equipment. This work was funded by RIKEN BSI and RIKEN CBS institutional funding, JSPS grants 26290011, 17H06037, and C0219129, and a Fujitsu collaborative grant (to A.B.); RIKEN JRA, University of Tokyo IST-RA, and JSPS-DC2 (to F.B.); HFSP postdoctoral fellowship LT000582/2019 (to J.O.); and SPDR (to R.A.).

## Data availability

Data will be deposited in: https://github.com/CBS-NCB/mouseTexturesData.

## Code availability

Analysis code is available at this GitHub repository: https://github.com/CBS-NCB/mouseTexturesCode.

## Author Contributions

A.B., J.G., F.B., and J.O. designed the study. A.V.J. facilitated image generation. R.A. helped design the protocol for behavioral training. F.B. created the behavioral protocol, collected all behavioral data, performed all recordings, and conducted all analyses. A.B. and F.B. wrote the manuscript. A.B. supervised the work.

## Materials and methods

### Subjects

All procedures were reviewed and approved by the Animal Care and Use Committees of the RIKEN Center for Brain Science. The behavioral data for the texture–scramble and texture–texture discrimination visual task were collected from a total of 21 mice: six CamktTA;TREGCaMP6s (four males and two females), 14 C57BL/6J WT (11 males, three females), and one male CaMKII*α*-Cre. For the passive widefield and two-photon imaging experiments, we used a total of 11 mice: six CaMKII*α*-Cre transgenic mice (four males and two females) and five C57BL/6J WT (two males and three females). The age of the animals typically ranged between eight and 28 weeks old from the beginning to the end of the experiments. Mice were housed under a 12–12 h light–dark cycle.

### Cranial window implantation

As described in Aoki et. al., 2017 [64], for the implantation of a head-post and optical chamber, the animals were anesthetized with gas anesthesia (Isoflurane 1.5–2.5%; Pfizer) and injected with an antibiotic (Baytrile, 0.5 ml, 2%; Bayer Yakuhin), a steroidal anti-inflammatory drug (Dexamethasone; Kyoritsu Seiyaku), an anti-edema agent (Glyceol, 100*µ*l; Chugai Pharmaceutical) to reduce brain swelling, and a painkiller (Lepetan, Otsuka Pharmaceutical). The scalp and periosteum were retracted, exposing the skull, and then a 5.5 mm diameter trephination was made with a micro drill (Meisinger LLC). Two 5 mm coverslips (120-170*µ*m thickness) were positioned in the center of the craniotomy in direct contact with the meninges, topped by a 6 mm diameter coverslip with the same thickness. When needed, Gelform (Pfizer) was applied around the 5 mm coverslip to stop any bleeding. The 6-mm coverslip was fixed to the bone with cyanoacrylic glue (Aron Alpha, Toagosei). A round metal chamber (7.1 mm diameter) combined with a head-post was centered on the craniotomy and cemented to the bone with dental adhesive (Super-Bond C&B, Sun Medical), which was mixed with a black dye for improved light absorbance during imaging.

### Viral injection

For imaging experiments, we injected the viral vector rAAV1-syn-jGCaMP7f-WPRE (4 × 1012 gc/ml, 1000 nl) into the mice’s right visual cortex (AP, −3.3 mm: LM 2.4 mm from the bregma) at a flow rate of 50 nl/min using a Nanoject II (Drummond Scientific, Broomall, Pennsylvania, USA). The injection depth was 400*µ*m. After confirmation of fluorescent protein expression (approximately two weeks after the AAV injection), we made a craniotomy (5.5 mm diameter) centered on the injection site while keeping the dura membrane intact and implanted a cover-glass window, as described above.

### Behavior

#### Behavioral training procedure

Mice were habituated to our automated behavioral training setups with self-head fixation, as previously described in ref [64]. The training of mice, from naïve to expert, progressed according to four stages with increasing difficulty, both procedural and perceptual, and with the fourth stage involving the final tasks described in the Results section (both the texture–scramble and texture–texture tasks). In the first stage, trial timing and stimulus properties were already set as in the final stage (Figure 1d). However, 1) the “go” stimuli were shown in 70% of the trials (instead of 50% in the fourth stage); 2) the minimum wheel rotation required to trigger a response was 5° instead of 45°; 3) the maximum wheel rotation that was allowed during the last second of the ITI was larger (20° instead of 5°); 4) the reward size was 8*µl* instead of 4 *µl*. During this training stage, the mice learned the association between wheel rotation and water reward contingent on the stimulus presentation on the screen. After they learned to rotate the wheel contingent to stimulus presentation in at least 80% of the trials for three consecutive sessions, they were moved to the second training stage, with the following changes: 1) the “go” stimuli were shown in 70% of the trials; 2) the wheel rotation angle to signal a response was increased to 15°; 3) the maximum wheel rotation allowed during the ITI was decreased to 5°; and 4) the water reward was lowered to 4 *µl*. After the mice reached at least 70% hits for three consecutive sessions, they were moved to the third training stage, in which the only change was an increase in the wheel rotation angle to 30° to signal a response. After reaching at least 70% hits for three consecutive sessions, the mice were moved to the fourth and final training stages with 50% hit trials. Most of the mice started the training with the honeycomb or scales texture/scramble family. Afterwards, we randomly selected the next family until all four families were successfully discriminated against the corresponding scrambles. A texture–scramble family discrimination was considered completed when the mouse had a *d*^′^ *>* 1 consistently over 10 consecutive sessions. The training details for the texture–texture task are described in the “Texture–texture task” section.

#### The texture–scramble task

Mice were trained in a go/no go texture–scramble discrimination task. Their self-head was fixed twice a day in a behavioral setup [64] connected to their home-cage, which comprised a self-latching stage, a rubber wheel with a quadrature encoder sensor to read the wheel’s position [91], a spout that dispensed water drops (4 *µl*), and a computer monitor positioned in front of the latching stage. Mice were required to rotate the toy wheel with their front paws contingent on a texture stimulus shown on the screen (the “hit” trials were rewarded with a water drop; the “false alarm” responses were discouraged by presenting a full-field flickering checkerboard pattern for 10 seconds; no feedback was given for “misses” and “correct rejects”). Regarding the temporal structure of the trial (shown in Figure 1d), a session began with an ITI with an isoluminant gray screen (with the same mean luminance level of the texture and scramble images). The ITIs lasted for four to six seconds chosen from a randomly uniform distribution. Mice had to refrain from rotating the wheel, with movements during a one-second period before the onset of the visual stimulus extending the ITI by one second. The stimuli had a 50% chance of being either a go stimulus (texture exemplar) or a no-go stimulus (scramble exemplar). The parameters of the stimuli matched those used in the imaging experiments: 100° in visual angle, with a raised cosine mask to reduce sharp edges (high-frequency components), and the texture family to be discriminated was kept constant during the entire session, randomly selecting the image to be displayed in each trial from a set of 20 exemplars. Following the stimulus presentation, the mice had two seconds to respond (response window). A wheel rotation was counted as a response if it exceeded 45°. After a hit trial, a water reward was given, which was followed by a one-second period, during which the stimulus remained visible on the screen, which then disappeared at the beginning of the ITI period with a randomized four to six second duration. In false-alarm trials, the stimulus disappeared after the wheel rotation, and a flickering checkerboard pattern (2 Hz) was displayed for 10 seconds followed by an ITI period. For miss trials, a new ITI began at the end of the two-second response window.

The session ended either when the mice received 400 *µl* of water or when the session’s duration reached 1800 seconds. To verify that the mice did not rely on “brute force” memorization to solve the task [66], in a subset of expert animals (n = 17), trained on all four texture–scramble family pairs, we introduced new sets of texture and scramble exemplars (20 each) and compared the performance of mice in the five sessions before and after the change in exemplars.

#### Texture–texture tas

Mice trained in the texture–texture go/no go task were both a subset of the mice trained in the texture–scramble (n = 14) and a new cohort of naïve mice (n = 2). If the mouse had been previously trained in the texture–scramble task expert, we simply changed the protocol so that a randomly chosen texture family (20 exemplars) was the new “go” stimuli and, similarly, another randomly chosen texture family (20 exemplars) was the new “no go” stimuli. Instead, for naïve mice, we trained them following the same training procedure described for the texture–scramble task but using exemplars from another randomly chosen texture family instead of scrambles.

### Image synthesis

#### Texture synthesis

As described in Ding et al., 2020, convolutional neural networks (CNNs) can be used to extract a compact representation of texture images by measuring the activation patterns of a CNN for a given texture. These activations are an over complete multi-scale representation [59, 92] that can be used to synthesize an arbitrary set of texture exemplars. Specifically, the first step for the synthesis of a novel texture exemplar relative to a reference texture (“target”, x) is to obtain a CNN parametrization of x—that is, its feature vector representation, *f* (*x*). This is done by concatenating the spatial means of the feature-map activations in each of the five VGG16 layers, which results in a feature vector of size 1,472:

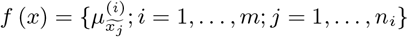

where *m* = 5 is the number of convolutional layers, *n*_*i*_ is the number of feature maps in convolutional layer *i*, 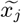 is the spatial mean across filter activations in the feature map j, and 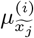 is the set of such means for layer i. The second step is to obtain a feature vector representation, *f* (*y*), of a Gaussian-noise image y:

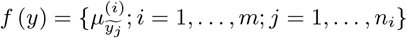

To obtain *f* (*x*) ≈ *f* (*y*), we solve an optimization problem (with an L1 loss):

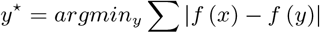

Where *y*^⋆^ is the fully optimized image relative to the target image, x. This approach is nearly identical to that of Ding et. al., 2020, with the only difference being that we did not add the mean of the three-color channels of *x* to our feature transform [59] since—in our framework—it created some degree of “pixelation” in the synthesized images. Rather, as an additional step after optimization, we normalized the images to have equal mean luminance and standard deviation (RMS contrast).

#### Texture normalization

To ensure that texture exemplars had the same lower-order statistics (mean luminance and RMS contrast), we z-scored the pixel intensity values, multiplied them by a fixed contrast (standard deviation, *σ* = 0.15), and, finally, added a fixed mean luminance value (*µ* = 0.5). This normalization was applied to all the “target” texture images (relative to the synthesis procedure with VGG16) and the synthetized texture exemplars, as there were small differences in the luminance and contrast relative to the target after each exemplar was synthetized. Furthermore, to ensure that the spatial frequency content of the textures was within the range of mouse perceptual sensitivity, we used an iterative algorithm in which we progressively rescaled the “target” texture images such that 1) *>* 95% of the spatial-frequency amplitudes of all the target textures lied within the 0.0 to 0.5 cpd [93] interval; 2) the average amplitude spectrum overlapped across families in the frequency range between 0.01 and 0.5 cpd.

#### Scramble generation

Scrambles are the noise images spectrally matched to the textures [32] generated via FFT-transform of a given texture exemplar (changing for different texture–scramble pairs) and randomizing the phase components while keeping the amplitude ones. Phase randomization was done by drawing the phase values from an FFT-transform of a Gaussian-noise image. The thus-generated scrambles retained the same average orientation and spatial-frequency power as the texture exemplars but lacked the higher-order statistical dependence of the textures [32]. For each of the synthesized scrambled images, we verified that the mean luminance and RMS contrast remained nearly identical to the original textures. The difference was within the floating point error.

### Image analysis

#### Image statistics

We explored the image statistics at various levels of complexity. Our texture normalization procedure ensured that the pixel histogram distributions had identical means and standard deviations between the images (i.e., luminance, and RMS contrast). Within families and between a matching pair of textures and scrambles, we also confirmed that the average orientation and spatial frequency content were the same. To do so, in each image Fourier transform, we measured the average power in “slices” of the spatial frequencies and orientations (spatial frequency bins: [0.01 cpd, step 0.02 cpd : 0.5 cpd]; orientation bins: [0° : step 15° : 180°]). The plots in Figure S1b,c show the amplitude values as a function of spatial frequencies and orientations, averaged across 20 exemplars for all families and stimulus types, and normalized to 1. To measure the higher-order statistics of the images, we decomposed them using an approach devised by Portilla and Simoncelli [14], which decomposes an image using a bank of linear and energy filters tuned to different orientations, spatial frequencies, and spatial positions. The correlations are then computed across the outputs of these filters (i.e., the “PS statistics”). The parameters and classification of the PS statistics we adopted follow what has been previously described [32, 36, 37, 40]. Briefly, we used a filter bank composed of four spatial scales (four downscaling octaves), four orientations (0º, 45º, 90º, 135º), and a spatial neighborhood of seven pixels to compute the filter output correlations. In addition, the marginal statistics of the pixel distributions were also computed (min, max, mean, standard deviation, skewness, and kurtosis). However, since part of our image synthesis pipeline procedure already ensured equal mean and standard deviation, only the differences in skewness and kurtosis were added to the characterization of the image statistics. In the end, the output of this image decomposition yielded four main groups of PS statistics: 1) marginal statistics (skewness and kurtosis); 2) spectral statistics; 3) linear cross-correlation statistics; and 4) energy cross-correlation statistics.

#### PCA of PS statistics

The number of parameters associated with the PS decomposition was relatively higher (740) than the total number of images (eight image categories—four texture families and four scrambles—and 20 exemplars per category with two rotations) leading to redundancies between the PS statistical components [36]. We thus reduced the number of parameters by applying PCA to each PS statistical group after z-scoring the parameter values. We retained at most eight components in each group, which explained at least 70% of the variance per group (Figure S5e). The marginal statistics with only two “dimensions” were excluded from this decomposition. After PCA, we again z-scored the outputs across exemplars to ensure that the range of parameter values between the groups of statistics was commensurate; this was necessary to gain interpretability of the distance metric later introduced, which was based on these reduced PS statistics. We also confirmed that the reduced PS statistics retained sufficient information to discriminate between textures and scrambles, with the energy cross-correlation statistics maximally distinguishing between them (Figure S6a,b).

### Imaging experiments

#### Visual stimuli

The visual stimuli were shown on a gamma-corrected monitor (widefield: IIYAMA Prolite LE4041UHS 40”, two-photon: IIYAMA Prolite B2776HDS-B1 27”). The size of the stimuli was always 100 degrees of visual angle with a raised cosine window to correct for sharp edges; the stimuli were shown in front of the mouse perpendicular to its midline, which pointed to the center of the screen. The animal was at a distance of ∼33 cm from the monitor for widefield experiments and ∼24 cm for two-photon experiments. For widefield recordings, the stimuli were presented for 250 ms, followed by 750 ms of an isoluminant gray screen (ITI) before a new trial started. Each mouse was shown 20 exemplars of four texture families and four scramble families (computed from the textures), a total of 10 times each exemplar, with 200 blank trials (i.e., trials with an isoluminant gray screen and no stimuli). This resulted in a total of 1600 trials with images and 200 trials with no stimulus (blanks). The presentation of each image/blank was fully randomized across the entire session. The two-photon experiments followed the same temporal structure as the widefield experiments; however, we reduced the number of repeats and added image rotations. Specifically, each mouse was shown 20 exemplars of four texture families and four scramble families: a total of eight times for each exemplar, and two rotations (0° and 90°) of each exemplar, with 160 blank trials. This resulted in a total of 2560 trials with images and 160 trials with no stimuli (blanks). We also recorded the responses to oriented gratings: 100 degrees in size, four orientations (0°, 45°, 90°, 135°), five spatial frequencies (0.02, 0.04, 0.1, 0.2, 0.5 cpd) and 15 repeats per stimulus.

#### Widefield imaging

As described in Orlandi et. al., 2021 [94], the awake mice were head-fixed and placed under a dual cube THT macroscope (Brainvision Inc.) for widefield imaging in tandem-lens epifluorescence configuration using two AF NIKKOR 50 mm f/1.4D lenses. We imaged the jGCaMP7f fluorescence signals using interleaved shutter-controlled blue and violet LEDs with a CMOS camera (PCO Edge 5.5) with an acquisition framerate of 60 Hz. This dual color recording method ensured that we could capture both the calcium-dependent GCaMP signal (blue light path) as well as the hemodynamic-dependent signal (violet light path), as previously reported in other studies[67]. The blue light path consisted of a 465 nm centered LED (LEX-2, Brainvision Inc.), a 475 nm bandpass filter (Edmund Optics BP 475 × 25 nm OD4 ø = 50 mm), and two dichroic mirrors with 506 and 458 nm cutoff frequencies, respectively (Semrock FF506-Di03 50 × 70 mm, FF458-DFi02 50 × 70 mm). The violet path consisted of a 405 nm centered LED (Thorlabs M405L2 and LEDD1B driver), a 425 nm bandpass filter (Edmund Options BP 425 × 25 mm OD4 ø = 25 mm), a collimator (Thorlabs COP5-A), and joined the blue LED path at the second dichroic mirror. The fluorescence light path traveled through the two dichroic mirrors (458 and 506 nm, respectively) and a 525 nm bandpass filter (Edmund Optics, BP 525 × 25 nm OD4 ø = 50 mm) and was finally captured with the PCO Edge 5.5 CMOS camera using the cameralink interface. Camera acquisition was synchronized to the LED illumination via a custom Arduino-controlled software. The frame exposure lasted 12 ms, starting 2 ms after opening each LED shutter to allow the LED illumination to stabilize.

#### Preprocessing the widefield data

Data preprocessing was done with custom Python and MATLAB code, with subsequent analyses done in Python. The continuously acquired imaging data were split into blue and violet channels. Then, as described in Orlandi et. al., 2021 [94], we corrected for the “hemodynamic component” by removing a calcium-independent component from the recorded signal. For every pixel, the blue and violet data were independently transformed into a relative fluorescence signal, Δ*F/F* = (*F* −*aF* −*b*)*/b*, where *F* is the original data, and the *a* and *b* coefficients are obtained by linear fitting each time series, i.e., *F* (*t*) ≈ *at* −*b*. Afterwards, for each pixel, the violet Δ*F/F* signal was low-pass filtered (6th order IIR filter with cutoff at 5 Hz) and linearly fitted to the blue Δ*F/F* signal: the hemodynamic-corrected Δ*F/F* signal was obtained as Δ*F/F* corr = Δ*F/F* blue − (*c*Δ*F/F* violet + *d*), where *c* and *d* are the coefficients from linearly fitting the low-pass filtered Δ*F/F* violet to the Δ*F/F* blue signal, i.e.,Δ*F/F* blue(*t*) ≈*c*Δ*F/F* violet(*t*) −*d*. The continuously acquired data was then split into trial periods comprising sequences of frames in a temporal window of [-500, +1000] ms relative to stimulus onset. This resulted in a tensor with seven dimensions: [stimulus type (texture or scramble), family type (4), exemplars (20), repeats (10), no. pixels X (256), no. pixels Y (230), no. frames]. Next, we averaged across repeats to obtain an “exemplar response tensor.”

#### Retinotopy maps

After the mice recovered from the cranial-window surgery (typically 3 to 4 days), we performed widefield imaging recordings during visual stimulation with moving gratings to obtain retinotopic. We used a standard frequency-based method [95] with slowly moving horizontal and vertical flickering bars and corrections for spherical projections [69]. Visual area segmentation was performed based on azimuth and elevation gradient inversions [68]. The retinotopic maps were derived under light anesthesia (Isoflurane) with the animal midline pointing to the right edge of the monitor (IIYAMA Prolite LE4041UHS 40”), centered relative to the monitor height, and with the animal’s left eye at ∼25 cm from the center of the screen.

#### Two-photon imaging

As described in Aoki et. al., 2017, the imaging experiments were performed using the two-photon imaging mode of the multiphoton confocal microscope (Model A1RMP, Nikon, Japan) with a Ti:sapphire laser (80 MHz, Coherent©, Chameleon Vision II). The microscope was controlled using the A1 software (Nikon). The objective was a 16x water immersion lens (NA, 0.8; working distance, 3 mm; Nikon). The field of view (512 × 512 pixels) was 532 *µm* × 532 *µm*. jG-Camp7f was excited at 920 nm, and the laser power was ∼40 mW. The images were acquired continuously at a 30 Hz frame rate using a resonant scanner. To align the two-photon imaging field of view with the retinotopy, we captured a vascular image at the surface of the cortex and used it for reference.

#### Preprocessing of two-photon data

All the analyses, except for neuronal segmentation, were conducted using a custom code written in Python. The cells were segmented using Suite2p [96], followed by the manual classification of the segmented ROIs. We then computed the Δ*F/F*_0_ response values (%) for each neuron by first applying a neuropil correction: *F*_c_–*F*_s_ −0.7*F*_*n*_, where *F*_c_ is the corrected signal, *F*_s_ is the soma fluorescence, and *F*_*n*_ is the neuropil fluorescence. Then, we computed a baseline-fluorescence value (*F*µ) as the mean of *F*_c_ during the first five seconds of the recordings when no stimuli were shown on the screen. We then detrended *F*_c_ (Scipy function scipy.signal.detrend) to remove the slow decrease in fluorescence sometimes observed across several tens of minutes and used the zero-mean detrended signal *F*_d_ to compute Δ*F/F*_0_ = *F*_d_*/F*_µ_.

## Data analysis: widefield

### Defining regions of interest

For every visual area, we defined a visually responsive ROI (or stimulus ROI) based on the maps of azimuth and elevation obtained from widefield imaging, so as to include a range of [+30°, −10°] in azimuth (relative to the contra- and ipsilateral visual fields, respectively) and elevation (±30°), which, for the azimuth, was a conservative estimate of the retinotopic representation of the stimuli (of size ±50° in azimuth and elevation).

### Peak-response and p-value maps

The widefield responses to textures and scrambles (Figure 2b) were computed by averaging across repeats, exemplars, and families; the frames were then averaged in a time widow [200, 400] ms after the stimulus onset, approximately centered around the time of peak response. The difference between these two images is shown in Figure 2c. The temporal response curves in V1 and LM to the textures and scrambles (Figure 2d) were computed by averaging across repeats, families, and pixels within the response ROIs in V1 and LM; the variability was across the exemplars. The response ROIs were defined based on retinotopy as the cortical region that “mapped” the stimulus location in the visual space. The error bands indicated a 95% confidence interval across the exemplars. To evaluate the significance of the differential response to the textures and scrambles (Figure 2e), we tested against a distribution of pre-stimulus responses. Specifically, we first computed the response–difference distributions by subtracting the responses to texture exemplars (averaged across repeats) from the randomly paired scramble exemplars. As before, the frames were also averaged around the time of the peak response, [200, 400] ms after the stimulus onset. This resulted in a tensor with four dimensions: [family type (4), exemplars (20), no. pixels X (256), no. pixels Y (230), no. frames]. By grouping the responses to all the families and exemplars, we generated response-difference distributions for each pixel, each containing 80 data points. We applied the same procedure to the data in a temporal window [-350, −100] ms prior to stimulus onset to obtain the “null” distributions for each pixel. Finally, we tested for statistical differences between the pre-and poststimulus onset distributions using a paired t-test and reporting the associated p-values. This procedure was applied to each animal, and the p-value maps were then used to compute the texture modulation of each visual area, as described below.

### Texture selectivity of visual areas

To determine how significantly a visual area was modulated by textures compared to scrambles (Figure 2f), we computed the proportion of the significantly modulated pixels (*p <* 0.01, from the p-value maps) within the stimulus ROI of each area (described in the section “Defining regions of interest”). This was separately computed in five visual areas (V1, LM, RL, AM, and PM) that were reliably segmented in all animals.

### Texture discriminability

To compute the texture–scramble discriminability values for V1 and LM (Figure 2g), we considered the responses to exemplars—separately for textures and scrambles—averaged across (i) repeats, (ii) pixels within stimulus ROIs (see section “Defining regions of interest”), and (iii) time frames within a window of [200, 400] ms after the stimulus onset. We then calculated a texture–scramble discriminability index (*d*^′^) in both brain areas as follows:

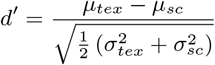

Where *µ*_tex_ and *µ*_sc_ are the mean responses to the texture and scramble exemplars (80), and 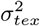 and, 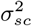 the corresponding variances. To calculate the “null distribution” of the *d*^′^ values shown in Figure 2g (gray band), we followed the same procedure as above in a time window [-300, −0] ms prior to the stimulus onset, reporting the 5% and 95% percentiles of that distribution.

## Data analysis: two-photon

### Stimulus-responsive cells

In a typical experiment, we could segment ∼200–450 cells (as described in the section “Two-photon imaging”). To establish whether a cell was visually responsive, in each trial ([-500, +1000] ms relative to stimulus onset) we “frame-zero” corrected Δ*F/F*_0_ by subtracting the average activity within a pre-stimulus period of [-500, 0] ms. Then, we used a *d*^′^ discriminability measure (similar to [69, 97]) by comparing the responses to visual stimuli and to “blanks.” Specifically, in each trial and for every segmented cell, we averaged the responses in a window of [250, 500] ms post stimulus onset. We then used these average values to generate two distributions: one from the trials with visual stimuli, the other from the “blank” trials. The distributions with the visual stimuli were computed separately for the individual texture and scramble exemplars and considering the response variability across repeats. For each stimulus exemplar, we then computed a discriminability measure, 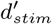, as follows:

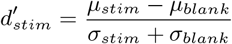

Where, *µ*_stim_ is the mean response across repeats for the chosen exemplar, *µ*_blank_ is the mean response across the repeats of blank trials, and *σ*_stim_, *σ*_blank_ are the corresponding standard deviations. This procedure generated a distribution of 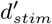 values for each cell. A cell was considered visually responsive if the maximum value of this distribution was ≥ 1, and if Δ*F/F*_0_ ≥ 6% in the stimulus-response window (for consistency with [69, 97]. Subsequent analyses were performed on this subset of stimulus-responsive cells.

### Texture–scramble d-prime

For every stimulus-responding cell, we considered frame-zero corrected Δ*F/F*_0_ data, averaging across repeats and responses in a time window of [250, 500] ms after stimulus onset. We then considered the data variability across exemplars (and their rotations) to compute a discriminability measure *d*^′^ as follows:

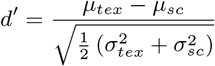

Where *µ*_tex_ and *µ*_sc_ are the mean responses to the texture and scramble exemplars, and 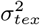 and 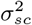 the corresponding variances.

### Regressive model

Using a set of reduced PS statistics as regressors (see section “Image statistics”), we constructed a linear regressive model (ridge regression) to predict individual cell responses. For each exemplar, we computed an average response value as the mean Δ*F/F*_0_ (averaged across repeats and frame-zero corrected) in a time window of [250, 500] ms post stimulus onset. For each neuron *i*, the model was trained to capture the responses to different exemplars using the following loss function:

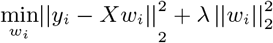

Where *w*_*i*_ are the optimization weights, *y*_*i*_ the data, *λ* a regularization parameter, and *X* the reduced PS statistics (two dimensions per group, i.e., the first two PCs). We confirmed that the model did not perform significantly better when using more PCs. The model was trained with five-fold cross validation to reduce overfitting, and the regularization parameter *λ* was optimized using a grid search. The model’s performance was evaluated in terms of the explained variance (EV) in the cross-validated data. To establish the significance of the model’s fit and to derive an EV threshold value for the inclusion of cells in the analyses of Figure 3h, we used a permutation test. For a given cell, we refitted the responses using as input statistics those from randomly chosen images (across exemplars from all textures and scrambles). Therefore, for each experiment, we obtained a shuffled distribution of EVs (across cells) and chose the 95th percentile of the distribution as the threshold value for significance (*α* = 0.05). We used this approach in all n = 20 experiments, resulting in an average threshold value, *EV*_*th*_ = 0.87% *±* 0.07% (s.e.). We set a conservative inclusion threshold at *EV*_*th*_ = 1%.

### Regressive model: weight analysis

To examine the contribution of the different reduced PS statistics in the regressive model, we summed the absolute values of the regressive weights separately for each of the four statistical groups: for a given cell, and for the PS group *i*, we computed 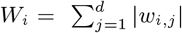, with *d* = 2, that is, the number of PCs for the reduced PS statistics. We then averaged across all the cells in a given animal (the individual data points in Figure 3h).

### Regressive model: unique EV

To examine the unique contribution to the explained variance by the different reduced PS statistics, we measured the loss in EV when training models without a particular statistical group. Specifically, considering a subset of cells with significant explained variance (EV *>* 10%), we first trained a model with all four groups of PS statistics (full model). Further, we trained four more models, each missing one of the four PS groups. We then computed a measure of unique variance explained, 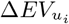, as follows:

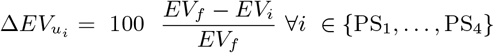

Where *EV*_*i*_ is the explained variance of a model trained without the PS group *i*, and *EV*_*f*_ is the explained variance of the full model.

### PCA embedding of neural response

For every stimulus-responding cell, we considered the frame-zero corrected Δ*F/F*_0_ data, averaging across repeats and time frames in a time window of [250, 500] ms post stimulus onset (as previously explained). After z-scoring the responses of each cell to different exemplars, we applied PCA (n = 20 PCs, separately for V1 and LM populations) to “standardize” the population size, thus facilitating a comparison between experiments, each having a different number of segmented cells. An example of a PCA space of neural activations is shown in Figure 4g for LM recordings (n = 2 PCs).

### Decoding responses to textures and scrambles

In the PCA spaces of neural activations for V1 and LM, as described in the section above, we considered responses to exemplars separately for each of the four texture–scramble families. For each family, we trained a binary logistic regression classifier to distinguish texture exemplars from scramble exemplars. The model was five-fold cross-validated, and its performance was evaluated using the average accuracy across the five folds. We repeated the same analysis by varying the number of PCs and examining the related changes in classification accuracy separately for the V1 and LM data (Figure S8a-c).

### Distance metrics for stimulus statistics

For each of the four PS statistical groups, we considered a 2D-PCA space of image statistics (see section “PCA of PS statistics”), with two PCs already sufficient for near-optimal classification performance (Figure S6a). The overall distance patterns described in Figure 4 were consistent when using larger numbers of PCs. A single point in each PCA space corresponds to the statistical representation of an exemplar image based on the associated PS statistical decomposition (reduced to four main PS statistical groups). To compute the radius of a cloud of points (20 exemplar points) for a given family, we computed the standard deviation of the x and y coordinates, *σ*_*x*_, *σ*_*y*_, and defined the radius as their mean value 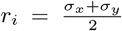. For the inter-cluster distance of a given pair of clouds (i.e., exemplars of textures or scrambles), we first computed the center of mass of the two clouds as the mean of the *x* and *y* coordinates, *µ*_*x*_, *µ*_*y*_, and then computed their Euclidean distance. The reported distance metric for each pair of the texture–scramble family was obtained by dividing the texture–scramble inter-cluster distance by the mean of the two corresponding radii. The inter-cluster distances were calculated for all the matching pairs of texture/scramble families (for Figure 4a-f). Further, the radius values were computed for all the families and stimulus types and for all the groups of PS statistics.

### Decoding the responses to texture families

In the PCA spaces of neural activations for V1 and LM (as described in the section “PCA embedding of neural responses”), we created another linear decoding model trained to classify all four texture families. We used a multinomial logistic regression classifier with an L1 regularization penalty. The training data consisted of the cells’ responses to 160 texture stimuli (four families, 20 exemplars, two rotations). The model was trained using five-fold cross validation, and the regularization factor was optimized with a grid search. The model’s performance was evaluated as the cross-validated accuracy averaged across folds. We also examined the dependence of the model’s performance on the number of PCs (Figure 4h).

### Distance metrics for neural representations

To compare the representational differences between V1 and LM, we created a common PCA space of neuronal activations. For a given mouse, we considered responses to exemplars pre-processed as described in “PCA embedding of neural responses” (before PCA). We then applied PCA to a “concatenated” ensemble of V1 and LM cells to derive a common PCA space with n = 8 components. The number of segmented cells and the z-scored response values were commensurate between V1 and LM. Using the PCA projection matrix, and by zeroing responses of the “other” area, we could then separately project the V1 and LM responses in this common space. We then measured the radii of the activation “clouds” in this PCA space for each texture family, as well as the inter-cluster distances for pairs of texture families, following the procedures described in “Distance metrics for stimulus statistics” (but without the normalization step for the inter-cluster distances). Finally, we compared the radii and inter-cluster distances for all six pairs of families between V1 and LM.

---

**Supplementary Fig. 1:**
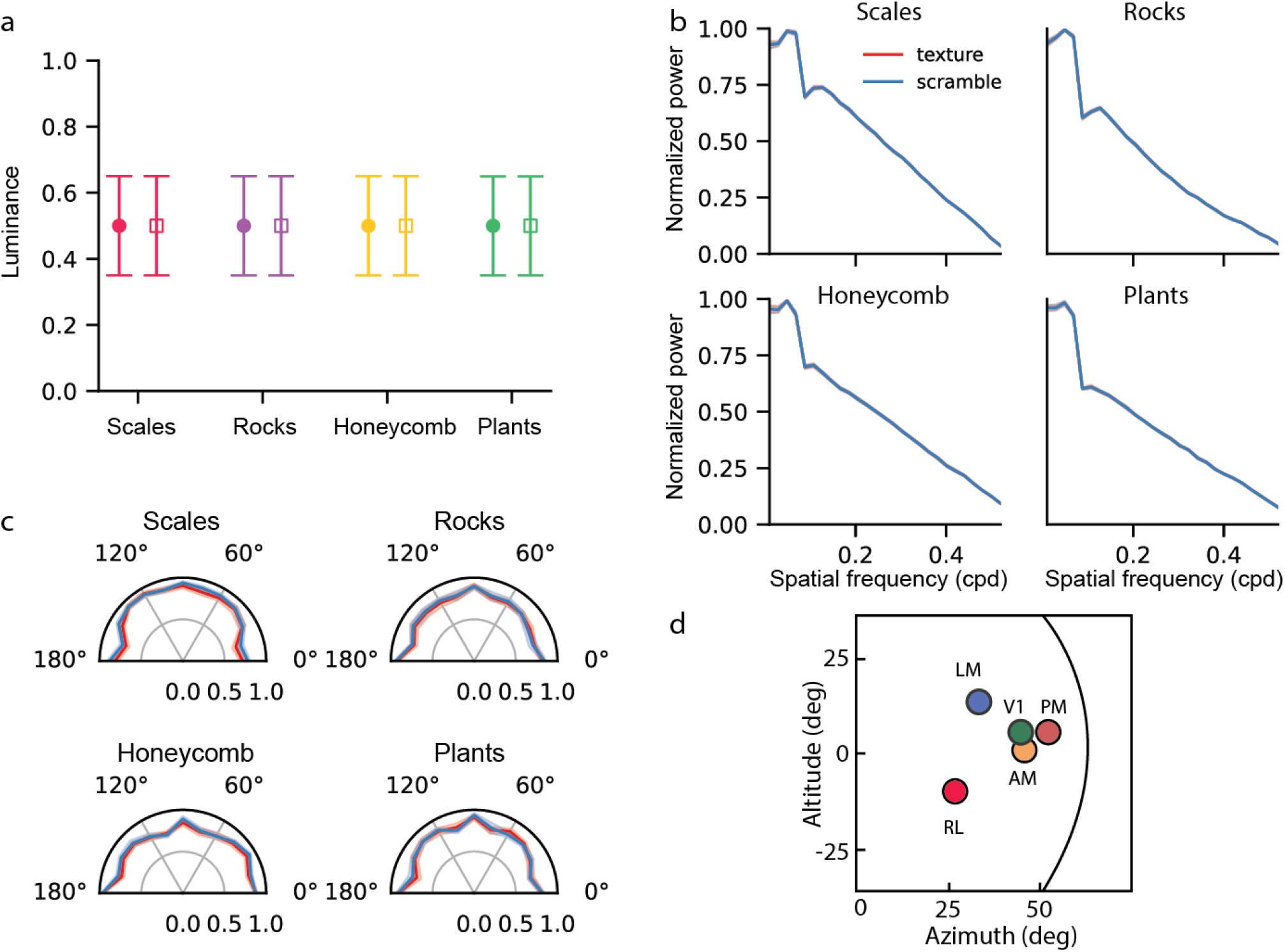
Characterization of the lower-order statistics of texture and scramble stimuli. **a**, The mean luminance across pixels for all texture exemplars (filled circles) and scramble exemplars (open squares) for each family (color code as in the main figures). The error bars indicate the RMS contrast–that is, the standard deviation of the pixel intensities averaged across all exemplars. **b**, The normalized spatial frequency power spectrum for each family—that is, the mean of the spectra computed for each exemplar, plotted in a frequency interval of maximum perceptual sensitivity for mice –0.02 - 0.5 cpd). **c**, The average orientation power for each texture family computed as the mean across exemplars for textures (red) and scrambles (blue). **d**, The mean azimuth and elevation of each reliably segmented visual area based on retinotopic mapping (filled dots, arbitrary colors; replotted from Garrett et al., J. Neurosci. 2014[68], Figure 6E); the semicircle delineates the size (fixed across experiments) of the visual stimuli.

**Supplementary Fig. 2:**
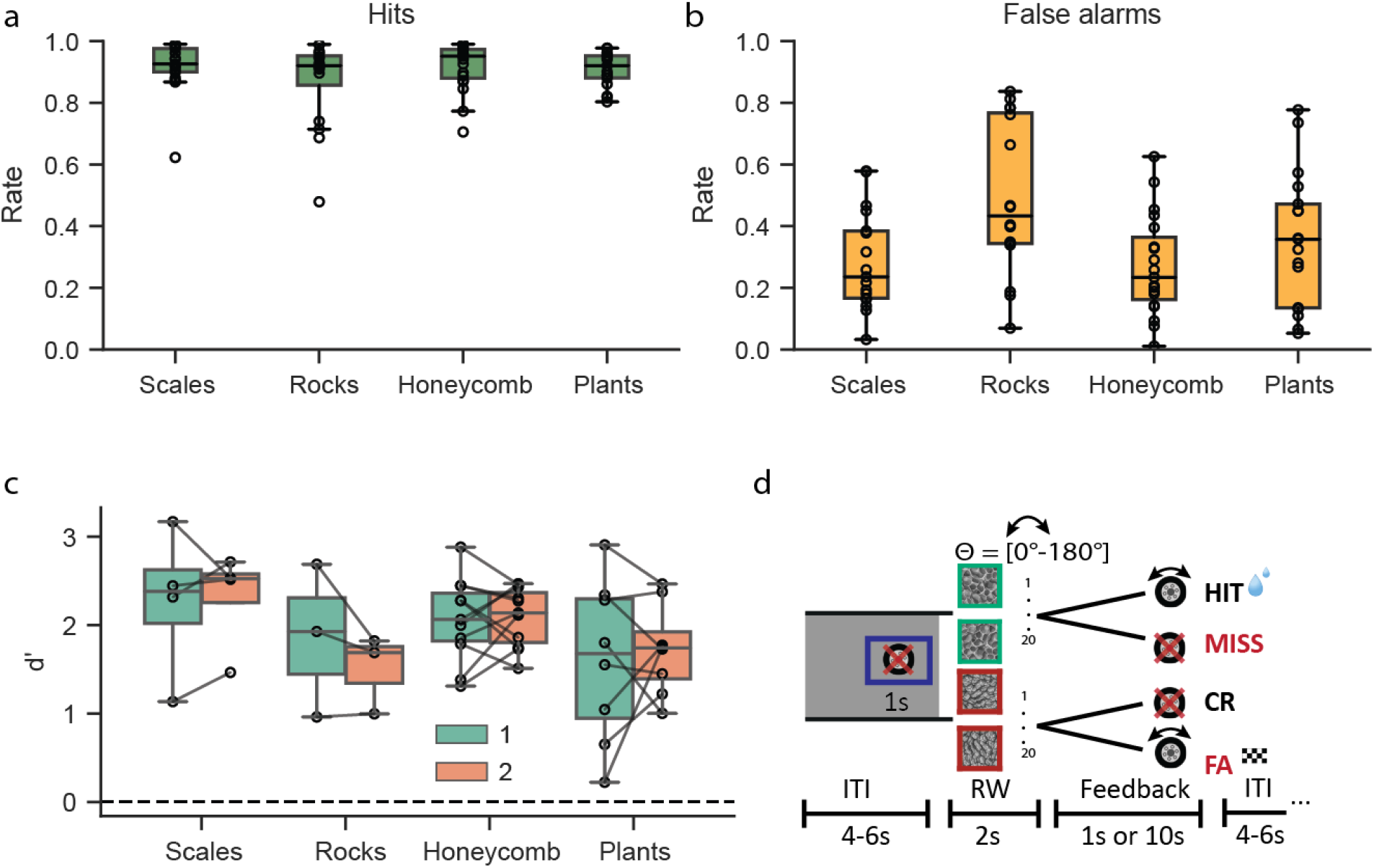
Performance of mice in the texture–scramble go/no go task. **a**, The average hit rates for each texture family for the mice shown in Figure 1f; each dot is for one animal. **b**, Same as in (a) but for false-alarm rates. **c**, The performance (*d*^′^) of mice (n = 3) for each texture–scramble pair over the last five sessions prior to a change in the set of 20 exemplars (green) and over the five sessions after the change (orange). Each dot indicates a training session; all differences are not significant. **d**, The texture/texture go no-go task: the mouse must rotate a wheel (go trial) if shown a go-texture exemplar; it must keep it still if shown an exemplar from the no-go family (no-go). ITI is the inter-trial interval, RW the response window, and the feedback period.

**Supplementary Fig. 3:**
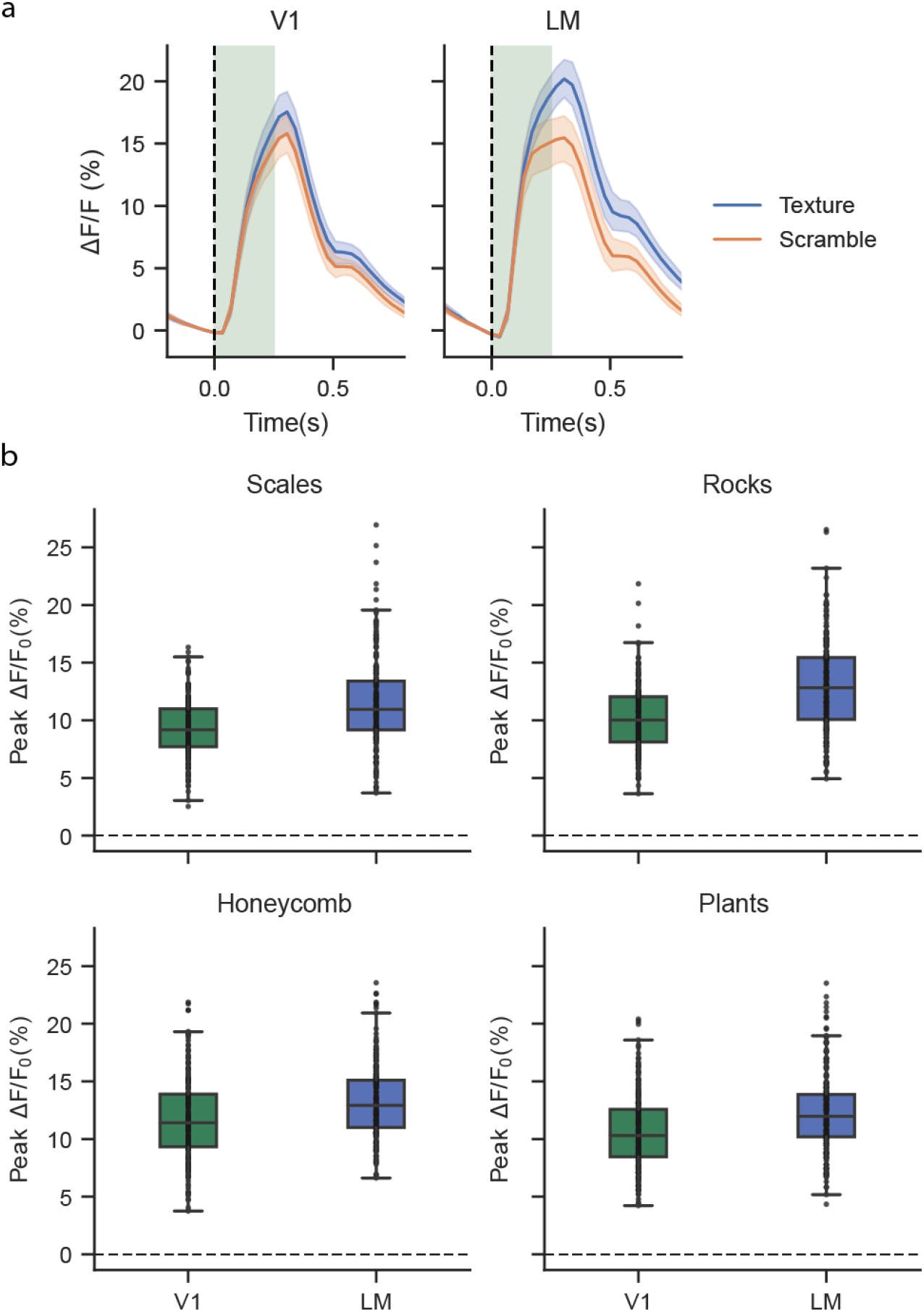
Population responses to visual stimuli derived from single-cell activity. **a**, The mean population activity of all the visually responsive cells per mouse (solid line, average across all mice; error bands, 95% confidence intervals across the average response per mouse (n = 10); Methods) as a measure of the collective population response to textures and scrambles for the V1 and LM experiments. The dotted vertical line indicates the stimulus onset, whereas the green band indicates the stimulus duration. The mean population activity was computed as the mean response to all the textures and scrambles, averaging across repeats, rotations, exemplars, and all the responsive cells. **b**, The population responses were calculated the same way as in (a), but the responses were separated across the families (4 plots), and only the texture stimuli responses were plotted for both V1 and LM.

**Supplementary Fig. 4:**
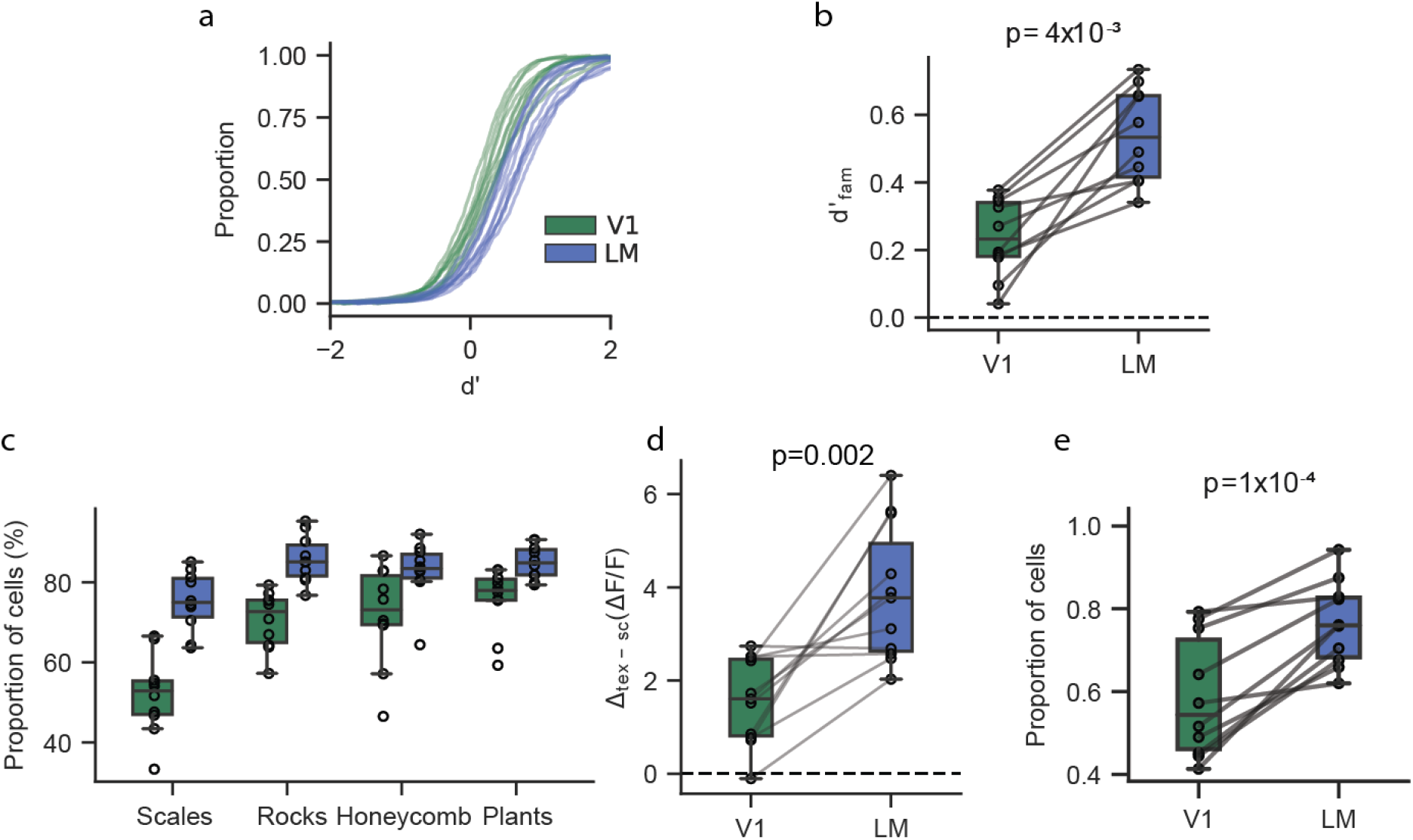
Single-cell responses to textures and scrambles. **a**, The cumulative distributions of the neural discriminability values (*d*^′^) of cells (combining across experiments and averaging across texture families) in the discrimination of textures vs scrambles images; green for V1 and blue for LM. Each line is for one animal. **b**, The average *d*^′^ values for each animal (black dots, n = 10 mice, mean across cells and texture–scramble pairs) for V1 and LM; the p-value from the paired t-test across mice (n = 10). **c**, The proportion of cells for each mouse (black dots) in V1 and LM with a *d*^′^ *>* 0 across all the texture–scramble pairs. **d**, The average modulation difference for each animal (black dots, n = 10 mice, mean across all cells, exemplars, and families) for V1 and LM; p-value = 0.002, paired t-test. **e**, the proportion of cells for each mouse (black dots) for which the regressive model based on the PS statistics had an explained variance EV ≥ 1%, separately for V1 and LM. The connecting lines are for the same animal; p-value, paired t-test.

**Supplementary Fig. 5:**
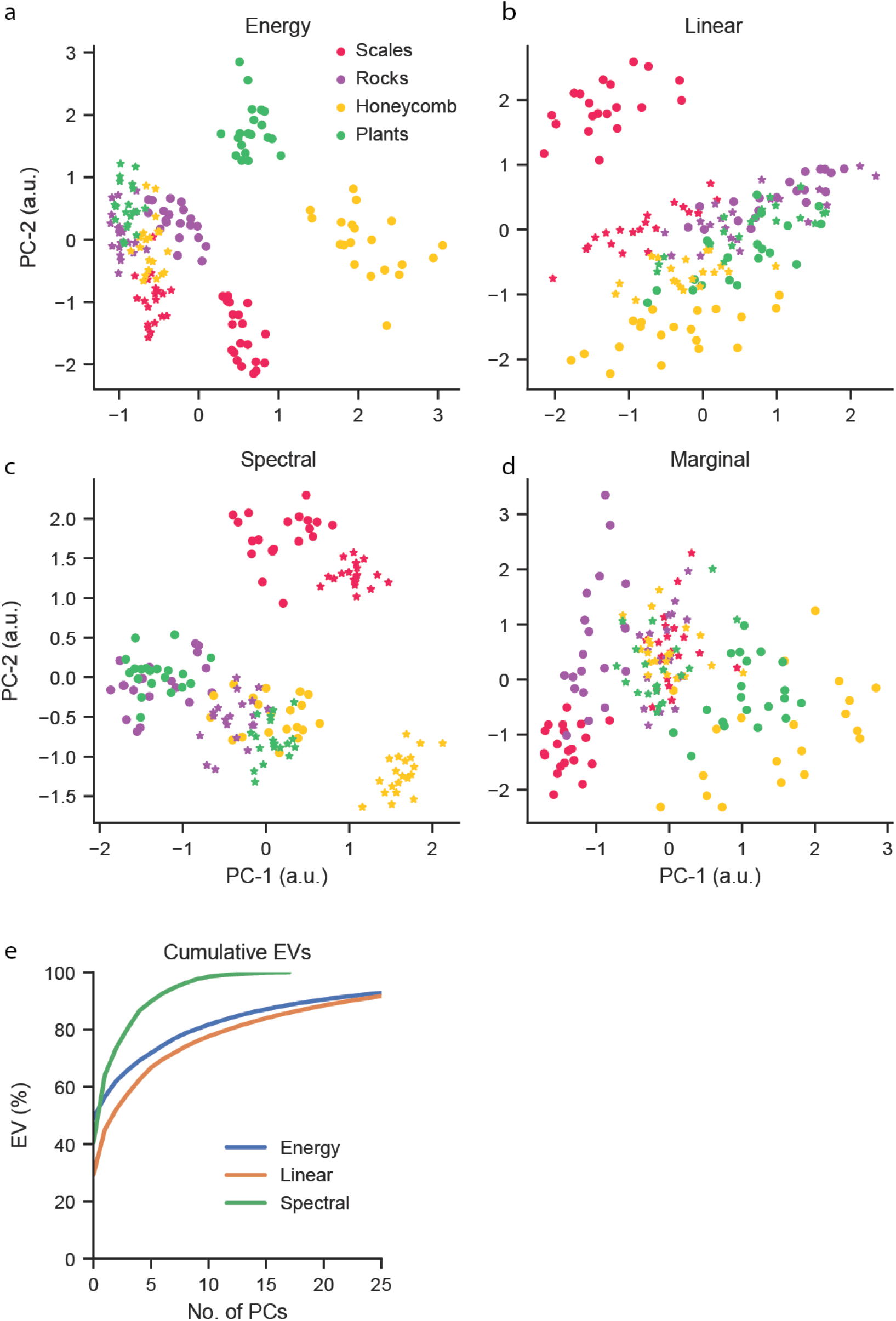
Clustering of visual stimuli in a PCA space of PS statistics. **a-d**,The two-dimensional PCA embedding of each of the four groups of image statistics (titles). The dots indicate (20) texture exemplars, and the stars (20) scramble exemplars. Color code for texture families in the legend. The same images were used for both behavioral and imaging experiments. **e**, The cumulative explained variance of the reduced PS statistics for the spectral, linear cross-correlation, and energy cross-correlation statistics.

**Supplementary Fig. 6:**
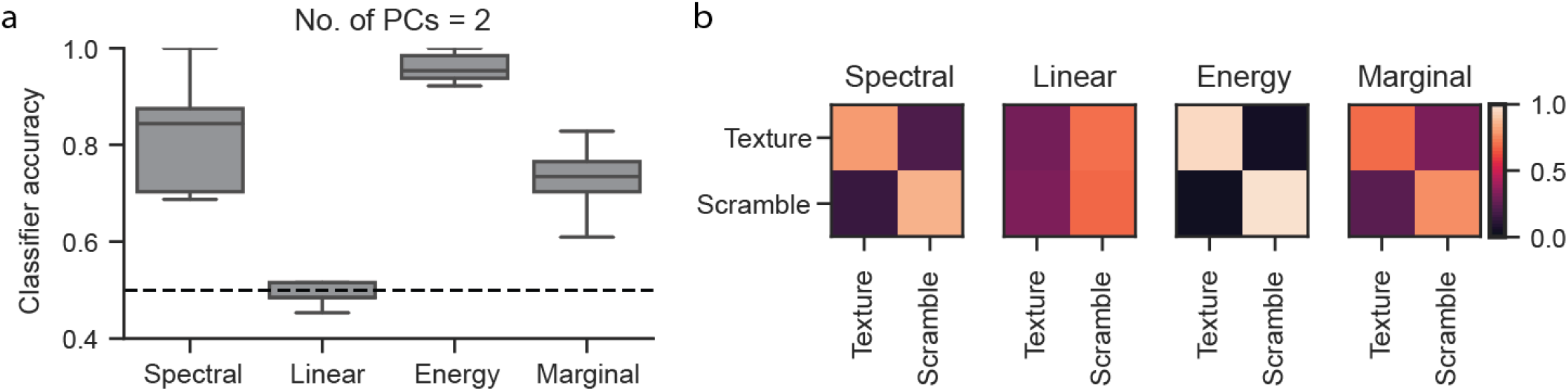
Binary texture–scramble classifier across a subset of PS image statistics. **a**, The cross-validated performance of a binary linear classifier trained to discriminate between texture and scramble images (across all families and exemplars) based on different PS statistical groups (x-labels). The horizontal dotted line indicates chance-level accuracy. The energy cross-correlation statistics is the group of image statistics with the highest discriminability accuracy in a 2D-PCA embedding space. **b**, The confusion matrix for the classifiers is shown in (a) for each PS statistical group (titles).

**Supplementary Fig. 7:**
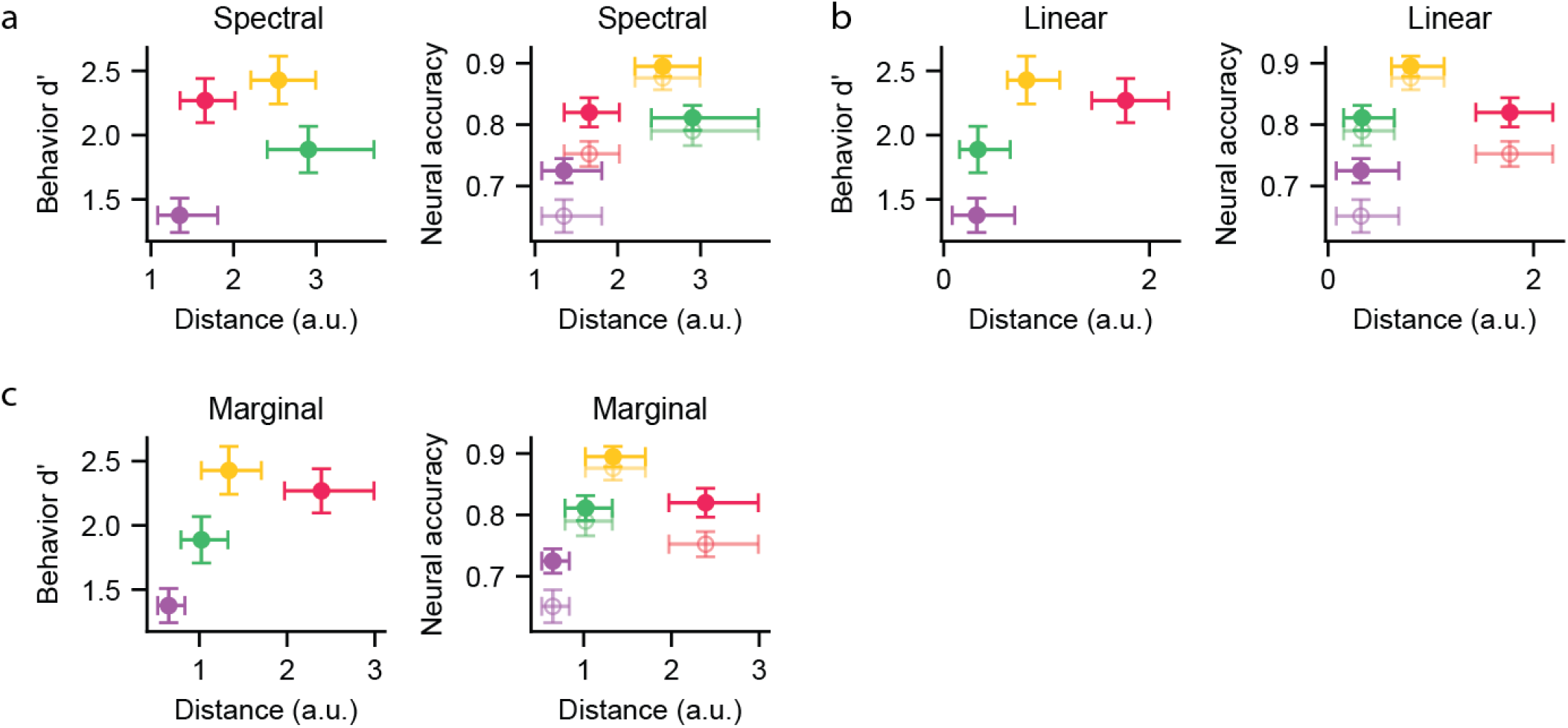
Neural, behavioral, and statistical distance measures for spectral, linear and marginal PS statistics. Each panel illustrates the same concept as in Figure 4, that is, the relationship between neural accuracy, behavioral performance, and image statistics, but for the other three groups of PS image statistics: spectral (**a**), linear cross-correlation (**b**), and marginal statistics (**c**). The error bars for behavioral performance and classification accuracy are the standard error of the mean; the error bars for inter-cluster distances are the 99.7% confidence intervals with Šidák correction for multiple comparisons.

**Supplementary Fig. 8:**
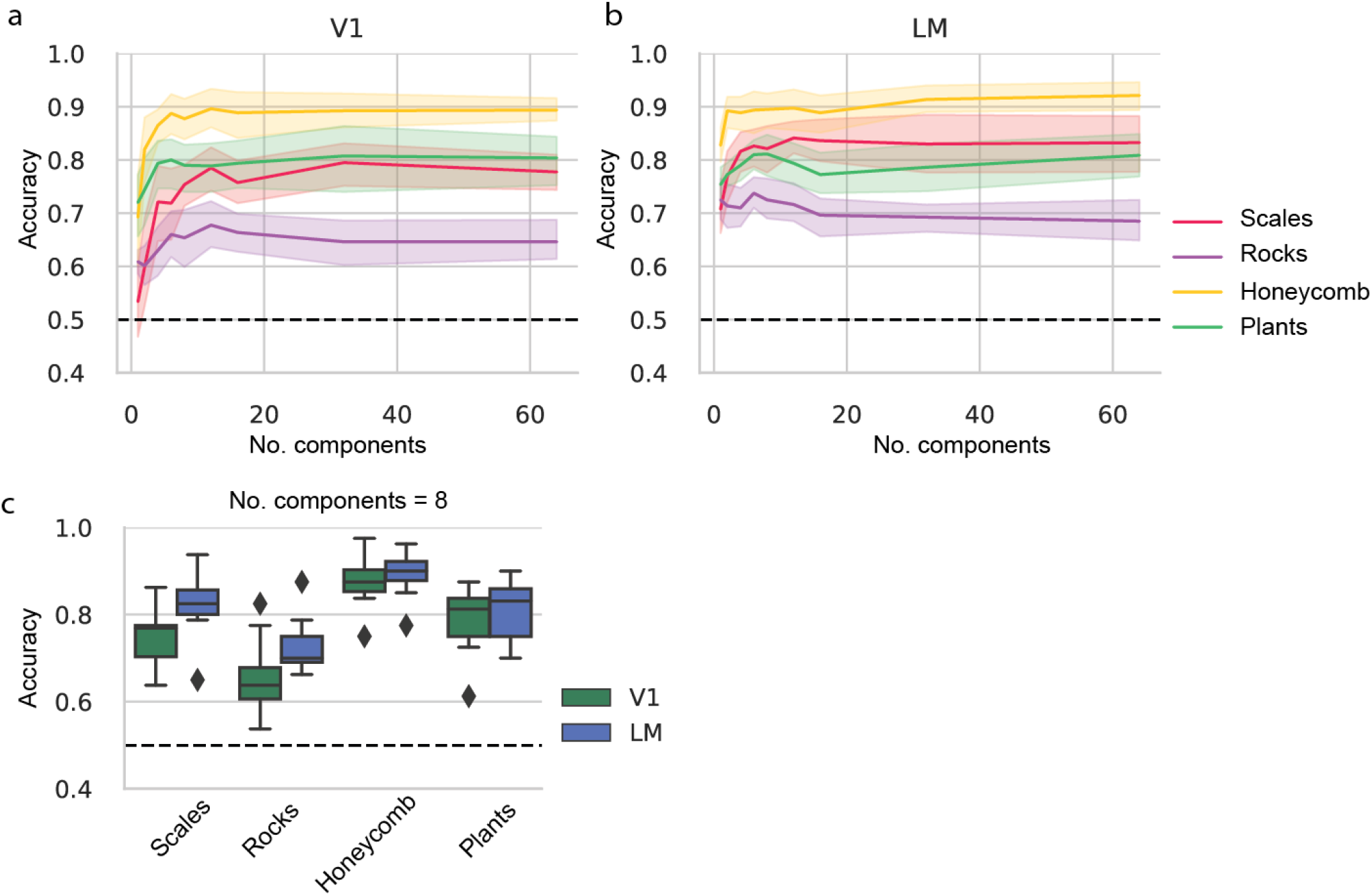
Binary classifiers of neural data for texture–scramble and texture–texture discrimination. **a**, The accuracy (fractional values) of a binary classifier trained on different pairs of texture–scramble families (legend in panel (b)) as a function of the number of components in the neural PCA space for V1. The shaded regions correspond to the 95% confidence interval for the average classification accuracy of all mice (n = 10). **b**, Same as in (a) but for LM. **c**, The accuracy of the same binary classifier in (a, b) when using eight PCA components.

**Supplementary Table 1:** Summary of the number of trials in the texture/scramble behavioral task.

| Mouse ID | Scales | Rocks | Honeycomb | Plants |
| --- | --- | --- | --- | --- |
| 20070 | 2189 | n/a | 1021 | 2029 |
| 20099 | 2015 | 1715 | 1122 | 2003 |
| 20100 | 2024 | 1979 | 1032 | 1677 |
| 20109 | 1853 | 1581 | 1000 | 1651 |
| 20117 | 1925 | 1938 | 1029 | 2213 |
| 21030 | 1471 | 1396 | 748 | 1682 |
| 21031 | 1816 | 1787 | 1056 | 1936 |
| 21032 | n/a | n/a | 1044 | n/a |
| 21033 | 1857 | 1560 | 1018 | 1856 |
| 21047 | 1839 | 1651 | 1985 | 756 |
| 21048 | 1002 | 1616 | 1852 | 1979 |
| 21049 | n/a | n/a | 871 | n/a |
| 21051 | 2059 | 1833 | 2049 | 870 |
| 21055 | 1666 | 1945 | 2059 | 864 |
| 21056 | 3712 | 1937 | 981 | 2040 |
| 21060 | 3845 | 2195 | 1118 | 2208 |
| 21061 | 1078 | 1892 | 2075 | 2172 |
| 21062 | 2042 | 2055 | 1962 | 1101 |
| 21064 | 3265 | 2332 | 2344 | 2260 |

**Supplementary Table 2:** Summary of the number of trials in the texture/texture behavioral task.

| Mouse ID | Honey/rocks | Honey/plants | Scales/plants | Rocks/plants | Honey/scales | Rocks/scales |
| --- | --- | --- | --- | --- | --- | --- |
| 20100 | 2097 | 2046 | 1587 | n/a | n/a | n/a |
| 20117 | n/a | 1637 | n/a | 1618 | 1583 | n/a |
| 21030 | n/a | n/a | 1971 | 2060 | 2038 | 2087 |
| 21031 | n/a | 1615 | 1918 | 2162 | 2144 | 1956 |
| 21033 | 1916 | n/a | 1775 | n/a | n/a | 2206 |
| 21047 | n/a | n/a | n/a | n/a | n/a | 1992 |
| 21048 | 1669 | n/a | n/a | n/a | 1786 | n/a |
| 21049 | 1890 | n/a | 2008 | 3727 | n/a | n/a |
| 21051 | n/a | n/a | n/a | n/a | 2091 | n/a |
| 21055 | 1676 | n/a | n/a | n/a | 1705 | n/a |
| 21056 | 1960 | 2022 | n/a | n/a | n/a | n/a |
| 21060 | 1802 | 1940 | n/a | n/a | n/a | n/a |
| 21062 | n/a | n/a | n/a | n/a | n/a | 1963 |
| 21064 | n/a | n/a | n/a | n/a | n/a | 2123 |
| 21067 | 1797 | n/a | n/a | 1619 | 1813 | n/a |
| 21074 | 2138 | n/a | n/a | n/a | n/a | n/a |

